# Simple biochemical features underlie transcriptional activation domain diversity and dynamic, fuzzy binding to Mediator

**DOI:** 10.1101/2020.12.18.423551

**Authors:** Adrian L. Sanborn, Benjamin T. Yeh, Jordan T. Feigerle, Cynthia V. Hao, Raphael J. L. Townshend, Erez Lieberman Aiden, Ron O. Dror, Roger D. Kornberg

## Abstract

Gene activator proteins comprise distinct DNA-binding and transcriptional activation domains (ADs). Because few ADs have been described, we tested domains tiling all yeast transcription factors for activation *in vivo* and identified 150 ADs. By mRNA display, we showed that 73% of ADs bound the Med15 subunit of Mediator, and that binding strength was correlated with activation. AD-Mediator interaction *in vitro* was unaffected by a large excess of free activator protein, pointing to a dynamic mechanism of interaction. Structural modeling showed that ADs interact with Med15 without shape complementarity (“fuzzy” binding). ADs shared no sequence motifs, but mutagenesis revealed biochemical and structural constraints. Finally, a neural network trained on AD sequences accurately predicted ADs in human proteins and in other yeast proteins, including chromosomal proteins and chromatin remodeling complexes. These findings solve the longstanding enigma of AD structure and function and provide a rationale for their role in biology.

## INTRODUCTION

Transcription factors (TFs) perform the last step in signal transduction pathways. They thus serve key roles in central processes such as growth, stress response, and development, and their mutation or misregulation underlies many human diseases (Spitz and Furlong, 2012). A TF includes a sequencespecific DNA-binding domain (DBD) and an effector domain that regulates nearby gene transcription. Activation domains (ADs) – effector domains that increase transcription – have long been of particular interest due to their roles as oncogenic drivers and use as scientific tools (Bradner et al., 2017; Brückner et al., 2009; Dominguez et al., 2016).

ADs were discovered as regions that could independently stimulate transcription when ectopically recruited to a gene promoter (Brent and Ptashne, 1985). Early experiments showed that ADs were unlike structured domains because progressive truncations showed graded reductions in activity (Hope and Struhl, 1986; Hope et al., 1988). Subsequent studies showed that ADs were disordered and had few similarities in their primary sequence (Mitchell and Tjian, 1989). Instead, ADs were classified based on their enrichment of certain residues, whether acidic, glutamine-rich, or proline-rich.

Acidic ADs are the most common and best characterized. Acidic ADs retain activity when transferred between yeast and animals, pointing to a conserved eukaryotic mechanism (Fischer et al., 1988; Struhl, 1988). While some have found that acidic residues are necessary for activation, others have found that they are dispensable (Brzovic et al., 2011; Pacheco et al., 2018; Staller et al., 2018; Warfield et al., 2014). Besides their negative charge, acidic ADs are rich in bulky hydrophobic residues. Mutating these hydrophobic residues reduces activation, often in proportion to the number mutated.

Because AD sequences are highly diverse and poorly conserved, only a small fraction of all ADs in eukaryotic TFs have likely been annotated. Sequence motifs have been proposed based on analysis of select ADs but have not been used for large-scale prediction (Piskacek et al., 2007). Screens of random sequences in yeast identified many activating sequences that represented as many as 1-4% of elements tested (Erijman et al., 2020; Ravarani et al., 2018). Actual protein sequences are, however, highly non-random. Direct screening of protein sequences has identified relatively few ADs at low resolution (Arnold et al., 2018; Tycko et al., 2020). There is a need for methods to experimentally detect or computationally predict all ADs.

ADs stimulate transcription of genes by recruiting coactivator complexes, especially the Mediator complex, which interacts with RNA Polymerase II and regulates its transcription initiation (Kornberg, 2005). Mediator is necessary for TF-dependent activation *in vitro* and *in vivo* and is required for regulation by enhancers in all eukaryotes (Allen and Taatjes, 2015). Mediator and TFs are concentrated at strong enhancers, perhaps in a phase-separated state, which may play a role in gene activation (Boija et al., 2018; Chong et al., 2018; Sabari et al., 2018; Shrinivas et al., 2019; Whyte et al., 2013). Beyond Mediator, TFs have been suggested to recruit other conserved multiprotein complexes, including TFIID, SAGA, and SWI/SNF, that play roles in activation of various genes (Hahn and Young, 2011; Mitchell and Tjian, 1989). While many such TF interactions have been described, their occurrence and roles remain to be determined.

Biochemical studies of TF-coactivator interaction have given insight into the mechanism of activation. Nuclear magnetic resonance (NMR) studies revealed short alpha helices formed by acidic ADs of yeast Gcn4 protein, with hydrophobic faces contacting hydrophobic surfaces of the Med15 subunit of Mediator (Brzovic et al., 2011; Tuttle et al., 2018). NMR constraints were consistent with multiple possible AD binding poses, suggesting a dynamic “fuzzy complex” (Tompa and Fuxreiter, 2008). Further consistent with these ideas, solvent exposure of aromatic residues was associated with activation in a screen of Gcn4 AD variants (Staller et al., 2018).

Here we combine quantitative, high-throughput measurements of *in vivo* activation and *in vitro* interaction with computational modeling to characterize ADs. We identify all ADs in budding yeast and describe their shared sequence attributes. We extend the analysis by training and validating a neural network that predicts new ADs across eukaryotes. Guided by predictions, we design and measure activation of thousands of AD mutants to derive a deeper understanding of the principles underlying activation. Correlating activation with measurements of binding of ADs to Mediator *in vitro* identifies the key protein interactions driving activation, for which we predict atomic structures using a peptide-docking algorithm. We derive structural features common to AD-Mediator interactions and use them to explain measurements of Mediator recruitment kinetics *in vitro.* In total, we develop an integrated model for function, sequence determinants, molecular interactions, and kinetic features of TF ADs.

## RESULTS

### A quantitative screen identifies 150 activation domains from all yeast transcription factors

We identified ADs across all TFs in budding yeast with the use of a quantitative, uniform, and high-throughput activation assay. Variability due to protein expression, activation duration, and secondary genetic effects was minimized by fusing domains of interest to a three-part artificial TF (aTF) that (1) is tracked by an mCherry tag, (2) localizes to the nucleus only upon induction with estrogen, and (3) binds uniquely through its mouse DBD in the promoter of a chromosomally-integrated GFP reporter gene (Figure 1A) (McIsaac et al., 2013; Staller et al., 2018). In this assay, three known ADs stimulated GFP expression greater than 100-fold upon estrogen treatment (Figure S1A).

**Figure 1.**
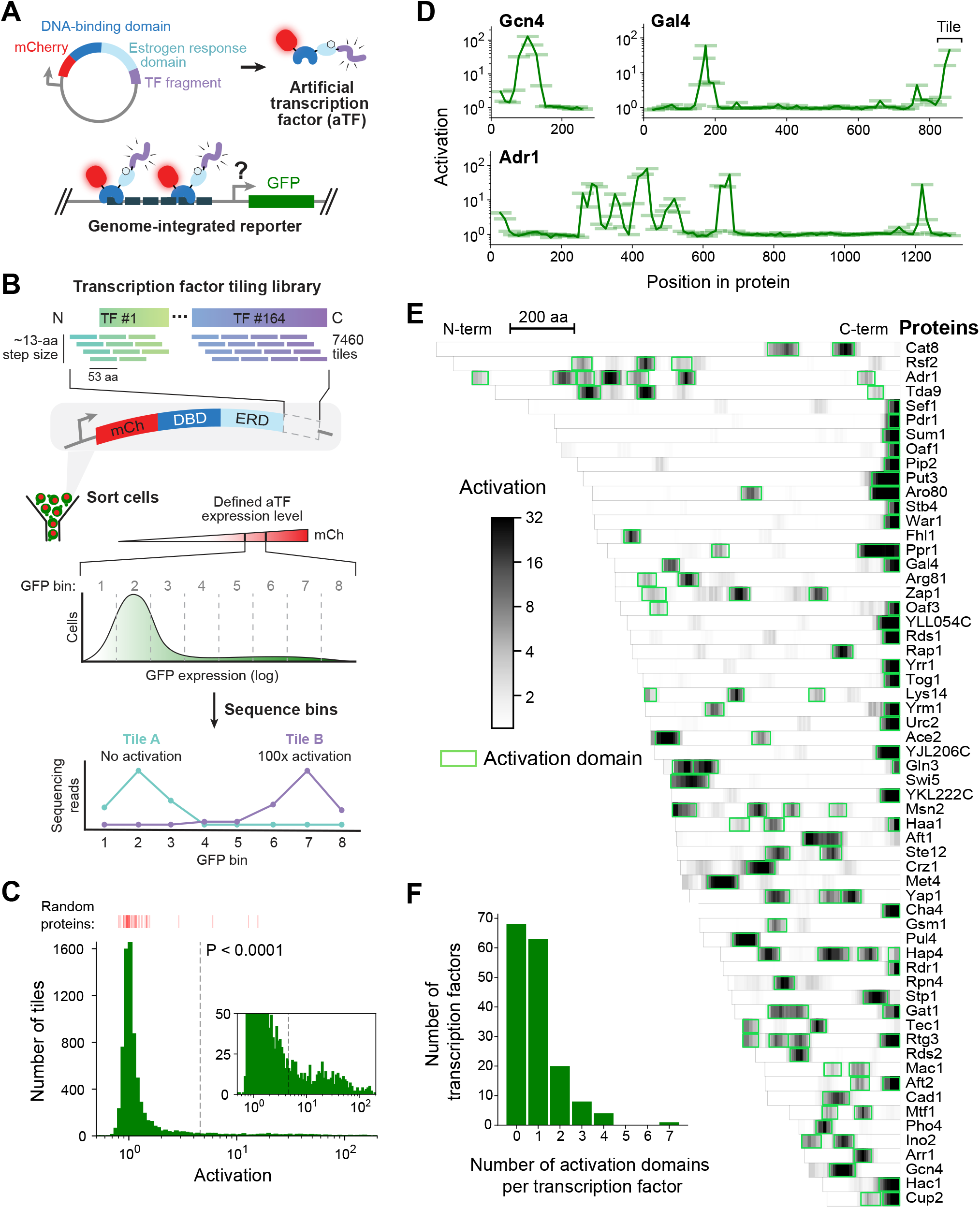
A quantitative screen identifies 150 activation domains from all yeast transcription factors. **(A)** Schematic of the activation assay. To measure *in vivo* activation, we expressed fragments of transcription factor (TF) proteins fused to a DNA-binding domain that binds uniquely in the promoter of a genome-integrated GFP reporter gene. This artificial TF (aTF) is tracked by its mCherry tag and localizes to the nucleus only after induction with estrogen. **(B)** Pooled screen for activation domains (ADs). All 164 yeast TFs were tiled by a DNA oligonucleotide library expressing 7460 protein segments, each 53 amino acids (aa) in length, with a step size of 12-13 aa (3.8-fold coverage). The library was cloned into the aTF expression plasmid and transformed into yeast cells. Using fluorescence-activated cell sorting, cells with a defined level of aTF expression were selected by mCherry signal and sorted into eight bins based on GFP expression. By next-generation sequencing, the distribution of each protein tile across the bins was determined and the mean value was used to calculate activation—namely, the fold increase in GFP relative to background. See also Figures S1A-D and Table S1. **(C)** Histogram of activation measured for 7460 tiles spanning all yeast TFs. Dashed line shows cutoff for P-values less than 0.0001 (Z-test). Red bars above the histogram mark activation of 50 random protein sequences. Inset: same histogram, zoomed in. See also Figure S1G. **(D)** Activation data for tiles spanning three example proteins. Activating tiles cluster into well-defined ADs when plotted by their protein position. Tiles, 53 aa long, are shown as horizontal bars with a line traced through their centers for clarity. See also Figure S1H. **(E)** Heatmap showing the mean activation at each position of the 60 TFs with the strongest ADs. Proteins run left to right from N-terminus to C-terminus and are sorted by length. A scale bar shows 200 amino acids (aa). ADs annotated in our screen are boxed in green and listed in Table S2. The method for annotating ADs is depicted in Figure S1H. **(F)** Histogram of the number of ADs in each TF. Of 164 TFs, 68 had no ADs, 63 had a single AD, and 33 had multiple distinct ADs, including up to seven distinct ADs in Adr1 (panel D). See also Figure S1I.

We used a pooled screen to measure activation by 7460 protein segments, each 53 amino acids (aa) in length, that tiled all 164 TFs with a step size of 12-13 aa (3.8-fold average coverage; Figure 1B). Outgrowth of transformed yeast for five days ensured that each cell contained a unique aTF expression plasmid due to mechanisms maintaining it at single-copy levels (Methods) (Scanlon et al., 2009). After induction with estrogen, cells with a defined level of aTF expression were selected by mCherry signal and sorted into eight bins based on GFP expression (Figure S1B). By nextgeneration sequencing, the distribution of each protein tile across the bins was determined and the mean value was used to calculate activation—namely, the fold increase in GFP relative to background (Table S1, Methods). Activation was highly concordant between distinct DNA sequences encoding the same protein fragment (Figure S1C). GFP distributions of three known ADs in the pooled screen exactly reproduced measurements from individual pure populations (Figure S1D). Controlling by a defined aTF expression level was critical for precisely measuring activation (Figures S1E-F). Activation measurements of fragments shared between two independently cloned and assayed libraries were reproducible (Pearson r = 0.953, Figure S1G). A median of 265 cells were assayed per tile and 99% of tiles were seen in at least 10 cells.

Our assay exhibited high signal-to-noise: tiles in previously known and newly identified ADs activated nearly 200-fold while 88% of fragments activated less than 2-fold (Figure 1C). Across the library, 451 tiles showed significant activation (P < 0.0001 by Z-test). When plotted by protein position, activating tiles clustered into discrete, well-defined ADs (Figure 1D). Using a positional activation score, we identified 150 ADs in 96 TFs (Figure 1E and S1H, Table S2, Methods). These ADs overlapped 75% of all previously-reported ADs in TFs (Table S2). Furthermore, the 53-aa tile length was not limiting, since our screen successfully identified ADs in over 85% of TFs that activated in a previous one-hybrid screen testing full-length proteins (Table S2) (Titz et al., 2006).

Three-quarters (112) of ADs identified here were previously unknown (Table S2). While 63 TFs contained just a single AD, 33 TFs had multiple, including up to seven distinct ADs in Adr1 (Figures 1D and 1F). C-terminal ADs were common—found in nearly half of all AD-containing TFs—and were stronger and shorter on average than other ADs (Figure S1I). Consistent with AD function in their native context, TFs that contained ADs upregulated a higher proportion of downstream genes than TFs without ADs (Figure S1J) (Hackett et al., 2020). These results show that our screen is both sensitive and comprehensive and has yielded the first complete annotation of ADs in any eukaryotic genome.

### Activation strength is primarily determined not by motifs but by acidic and hydrophobic content

ADs were enriched in negative charge and hydrophobic content, and activation was strongest for fragments that were high in both (Figures 2A-B and S2A; Wimley-White hydrophobicity scale) (Wimley and White, 1996). Past experiments have clearly shown the importance of bulky hydrophobic residues but presented conflicting evidence for acidic residues in activation (Brzovic et al., 2011; Pacheco et al., 2018; Staller et al., 2018; Warfield et al., 2014). To determine the role of negative charge, we took the strongest-activating fragment from each AD and measured its activation with all acidic residues mutated. Except for a single example that activated just as strongly at +7 net charge without its acidic residues (the Gln3 C-terminal AD), activation was abolished in all ADs, showing definitively that acidic residues are necessary in budding yeast ADs (Figure 2C). We also noticed that Asp but not Glu was associated with activation of wild-type tiles and hypothesized that Asp promotes activation more strongly than Glu (Figure S2C). Indeed, ADs with all Glu residues mutated to Asp consistently activated as well as or better than ADs with all Asp residues mutated to Glu (Figure S2D).

**Figure 2.**
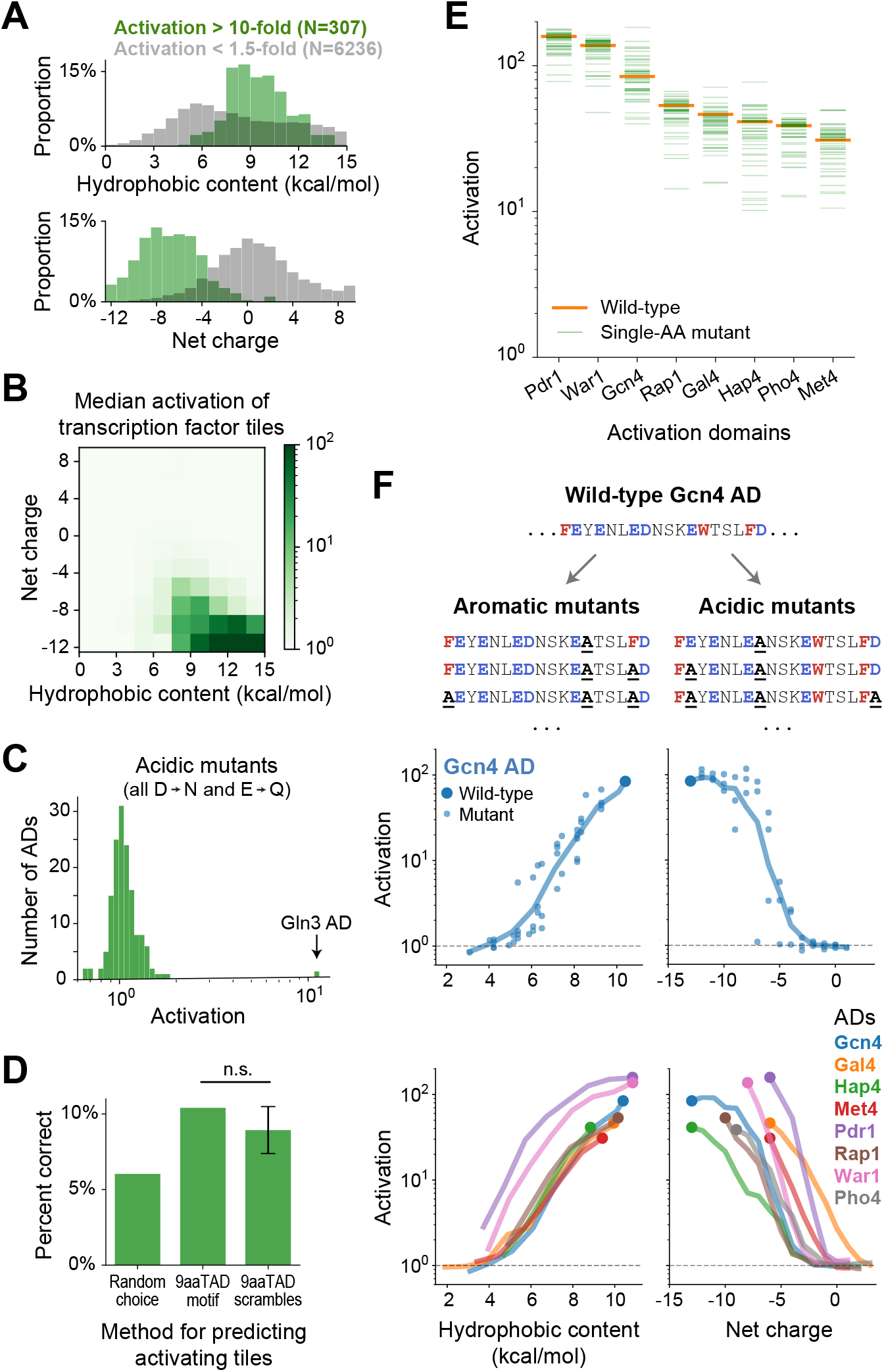
Activation strength is primarily determined not by motifs but by acidic and hydrophobic content. **(A)** Distribution of hydrophobic content (top) and net charge (bottom) for non-activating tiles (grey) and highly-activating tiles (green). Hydrophobicity is computed using the Wimley-White scale (Figure S2A) and charge is computed by counting Asp, Glu, Arg, and Lys residues. **(B)** Tiles are binned by their hydrophobic content and net charge and the median activation of each bin is displayed in color. Activation is strongest for tiles high in both acidic and hydrophobic content. **(C)** From each AD, activation of the strongest-activating fragment with all acidic residues mutated (Asp to Asn and Glu to Gln). Except for the Gln3 C-terminal AD, which activated just as strongly at +7 net charge without its acidic residues, activation was abolished in all ADs. See also Figures S2B-D. **(D)** Just 10% of tiles containing the 9aaTAD motif activated, which was only 1.7-fold better than guessing activating tiles at random and not significantly more predictive than scrambled versions of the motif. (n.s., not significant.) **(E)** Activation of all single amino-acid mutants (to Ala) across eight ADs. All mutants still activated more than 10-fold, and 94% of mutations affected activation by less than 2-fold. Activation for all mutants of two ADs are plotted by position in Figure S2E. **(F)** We varied the acidic and hydrophobic content of eight ADs by mutating successively larger subsets of aromatic (left) or acidic (right) residues. (*Top*) Example mutant sequences for a segment of the Gcn4 AD, with activation of the wild-type (large dot) and mutants (small dots) plotted below as a function of hydrophobic content or net charge. Lines trace a moving average of activation. *(Bottom)* Average activation as a function of hydrophobic content or net charge, for all eight ADs tested. Activation of all individual mutants is shown in Figure S2F. The Pho4 AD contains only two aromatic residues so its aromatic mutants are not shown.

Having identified the important residue types, we looked for sequence motifs that could predict activation. However, AD sequences appeared to share no common motifs upon inspection. The 9aaTAD motif, originally proposed based on yeast ADs (Piskacek et al., 2007), predicted ADs poorly: only 10% of 9aaTAD-containing tiles activated in our assay, which was not significantly more predictive than scrambled versions of the motif (Figure 2D). A *de novo* search for motifs using DREME also found none that were shared by more than two ADs (Bailey, 2011). Evidently, ADs are diverse in sequence and not predictable from simple motifs.

The lack of motifs suggests that activation is determined not by positions of individual residues but by distributed or redundant features. We pursued this point by measuring activation of a panel of designed mutants of eight ADs. Indeed, 94% of scanning single-amino acid mutants affected activation by less than 2-fold, and all still activated more than 10-fold (Figures 2E and S2E). To produce larger effects, we varied the acidic and hydrophobic content of the ADs by mutating successively larger subsets of acidic or aromatic residues (Figure 2F, Methods). Activation was gradually, and finally completely, eliminated as either characteristic was reduced. It is noteworthy that the Gcn4 AD, a focus of many past studies, uniquely tolerated mutation of up to six acidic residues. Most notably, activation by acidic and hydrophobic mutants were related to the number, rather than position, of mutated residues (Figures 2F and S2F). Thus, activation strength is primarily determined not by motifs but by acidic and hydrophobic content.

### A deep learning model, termed PADDLE, predicts the location and strength of acidic activation domains in yeast and human

Since motifs predicted ADs poorly, we turned to more sophisticated approaches using machine learning. We first trained a simple neural network to predict activation based solely on tiles’ amino acid (aa) composition (Methods). When tested on TF tiles withheld from training, this neural network performed remarkably well, explaining 66% of observed variation (Figure S3A). However, this algorithm was limited by its lack of information about residue positions. To experimentally disentangle the contributions of aa positioning and aa composition, we measured activation of eight scrambled sequences from each of the eight ADs tested before. Despite their shared abundance of acidic and hydrophobic residues, these mutants spanned a wide range of activation strengths, and some mutants activated stronger than wild-type while others failed to activate entirely (Figure 3A). Clearly, full sequence information beyond aa composition is needed to predict activation.

**Figure 3.**
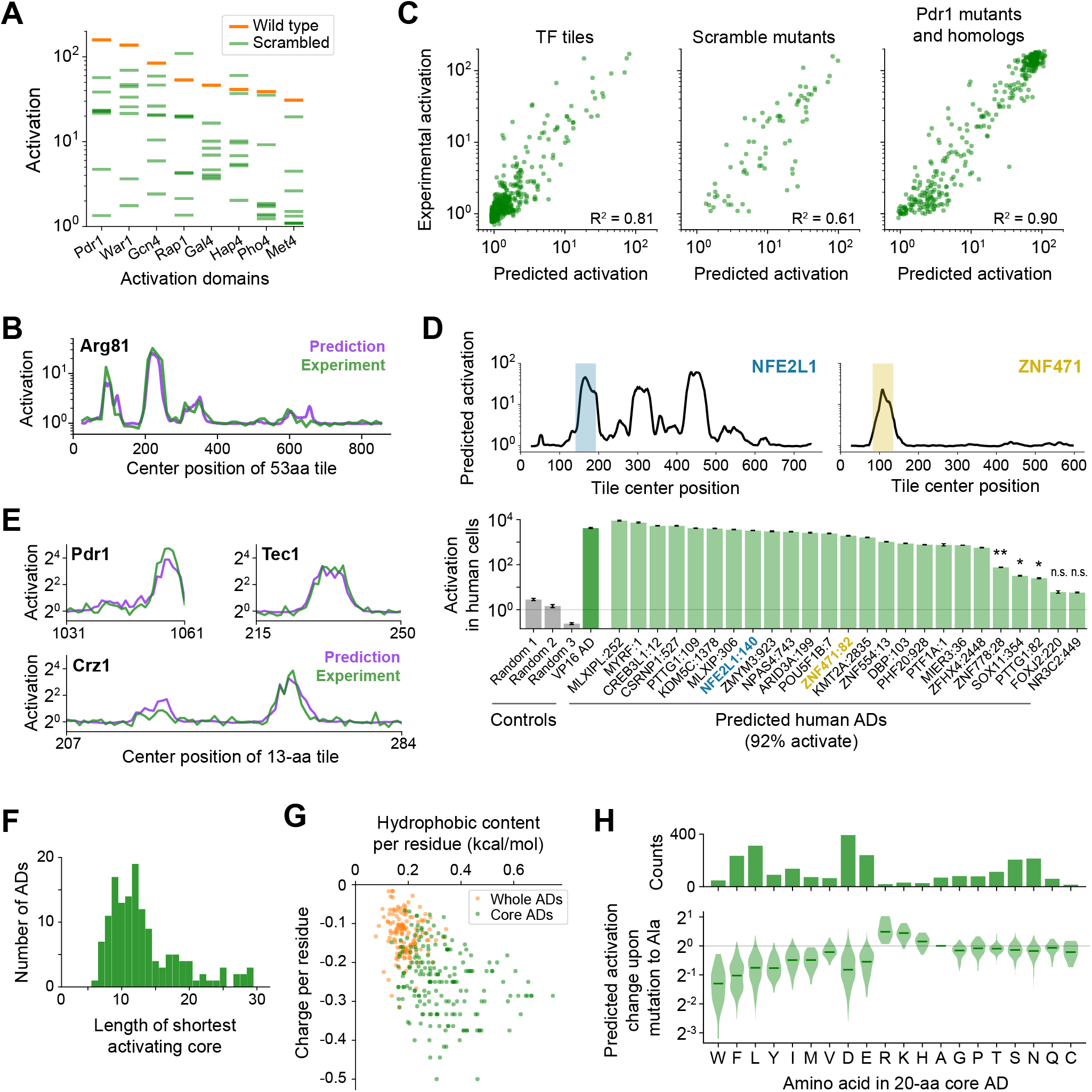
A deep learning model, termed PADDLE, predicts the location and strength of acidic activation domains in yeast and human. **(A)** Activation of wild-type (orange) and eight scrambled sequences (green) for each of eight ADs. **(B)** PADDLE predictions (purple) and experimentally-measured activation (green) for Arg81 are plotted by protein position. Predictions were run on 53-aa tiles in 1-aa steps and smoothed with a 9-aa moving average. **(C)** PADDLE predictions versus experimentally-measured activation is plotted for three categories of sequences that were omitted from PADDLE’s training dataset: wild-type tiles of TF proteins (left), the scrambled sequences of eight ADs from panel A (middle), and 232 mutants and 178 orthologs of the Pdr1 AD (right). R^2^ is the coefficient of determination. See also Figures S3A-C and Table S3. **(D)** PADDLE accurately predicts human ADs. (*Top*) Example PADDLE predictions on 53-aa tiles spanning two human TFs. One predicted AD from each TF, marked by the colored shading, was tested experimentally. *(Bottom)* We randomly selected 25 high-strength predicted ADs from human TFs and measured their activation individually using a luciferase reporter in HEK293T cells. Relative to three random sequence controls, 23 domains (92%) activated luciferase expression. The VP16 AD was used as a positive control. Error bars show standard deviation of technical triplicates. See also Figure S3D and Table S3. (* P < 0.01; ** P < 0.0005; n.s., not significant.) **(E)** PADDLE predictions (purple) and experimentally-measured activation (green) of 13-aa tiles in 1-aa steps spanning three ADs. Both predictions and experiments identified at least one significantly activating 13-aa tile within each of 10 ADs (Figures S3E-F). **(F)** PADDLE was used to identify the shortest core region of each AD that can independently activate. A histogram of their lengths is plotted. The minimal region for activation can be localized to within 20-aa core in 85% of ADs. **(G)** The hydrophobic content and charge per residue of whole ADs (orange) and core ADs (green). **(H)** Amino acid contributions to the strongest-activating 20-aa core within every AD. (*Top*) The number of times each amino acid is present. (*Bottom*) For each 20-aa core AD, the fold-change in activation upon mutating each individual residue to Ala was predicted using PADDLE. Those effects, grouped by the amino acid mutated, are shown in this violin plot. Median values are depicted by green lines. Mutants for three example core ADs are shown in Figure S3H.

We therefore trained a deep convolutional neural network (CNN) to predict activation based on protein sequence, predicted secondary structure, and predicted disorder (Methods). CNNs evaluate sequences by hierarchically integrating matches to a suite of learned patterns and have recently found great success in many genomic prediction tasks (Ching et al., 2018; Kelley et al., 2016; Wang et al., 2016). Our Predictor of Activation Domains using Deep Learning in Eukaryotes, or “PADDLE”, explained 81% of observed variation in TF tiles withheld from training (alternatively, area under the precision-recall curve of 0.805; Figures S3B-C, Table S3), markedly better than the aa composition-based predictor. *De novo,* PADDLE accurately predicted the activation strength of (1) new ADs within TFs omitted from training (Figures 3B-C, left); (2) scrambled AD sequences, despite their identical amino acid composition (Figure 3C, center); and (3) 232 mutants and 178 orthologs of the Pdr1 AD (Figure 3C, right). Altogether, the performance of PADDLE was validated across a wide range of both wildtype and mutant sequences in yeast.

Classic experiments showed that many acidic ADs retained activity even when transferred between yeast and animals (Fischer et al., 1988; Struhl, 1988), so we investigated whether PADDLE could identify ADs in human TFs that would activate in human cells. Using PADDLE, we predicted 236 high-strength and 366 moderate-strength ADs, together spanning 462 (27%) human TFs (Table S3). These predicted ADs overlapped many known ADs of TFs from diverse families, including p53, NFkB, Myc, Klf4, Fos, PPARA, SREBF1, E2F proteins, and the glucocorticoid receptor (Figure S3D).

PADDLE also predicted 41 high-strength and 45 moderatestrength ADs from among 419 transcription-regulating viral proteins (Table S3) (Liu et al., 2020). We randomly selected 25 high-strength predicted ADs from human TFs and measured their activation individually using a luciferase reporter in HEK293T cells. Remarkably, 23 domains (92%) activated luciferase expression (Figure 3D). Evidently, acidic activation mechanisms are conserved and the patterns learned by PADDLE are generalizable across eukaryotes.

### PADDLE identifies the core regions and key residues of every activation domain

To uncover the principles of acidic activation learned by PADDLE, we analyzed predictions on designed mutant sequences and developed hypotheses, which we then experimentally tested in a second library. We noticed from predictions on sequences less than 30 aa that yeast ADs contained short, independently activating regions, which we term core ADs (cADs) (Figure S3E). As an experimental test, we measured activation *in vivo* of 13-aa-long fragments tiling ten ADs in 1-aa steps (Figure 3E). Predictions proved highly accurate (R^2^ score = 0.842, Figure S3F), and both predictions and experiments identified at least one significantly activating 13-aa cAD within each AD (P < 0.001).

Across all yeast TFs, PADDLE predicted that the minimal region for activation could be localized to within a 20-aa cAD in 85% of ADs (Figure 3F, Table S3). These cADs still shared no motifs (Figure S3G), but instead had an especially high density of acidic and hydrophobic residues, even more so than the full ADs (Figure 3G). To quantify the contribution of each residue to activation, we predicted the impact of mutating each position in each cAD to alanine (Figure S3H). Single-aa mutants in these short cADs had a large impact on activation, unlike in the longer 53-aa ADs (Figure 2E), and showed clear trends when grouped by residue identity (Figure 3H). Mutating the bulkiest hydrophobic residues led to the greatest decreases in activation, and the three most important were Trp, Phe, and Leu. Mutating acidic or basic residues had opposite but comparable effects on activation. Together, these analyses identify the regions and residues across all ADs that contribute most to activation.

### A high density of acidic and hydrophobic residues is sufficient for most core ADs to activate, but some require a defined sequence and alpha helical structure

Having identified the core regions of every AD, we sought to understand the sequence properties that drive their activation. We focused on the 28 strongest 13-aa cADs (Table S3), and first quantified the importance of their aa composition by comparing each cAD with 33 random scrambles of its sequence. Remarkably, only nine cADs (32%) showed activation by the wild-type sequence greater than 3-fold greater than the average activation by scrambled sequences (Figure 4A). In these nine cADs, maximal activation evidently requires a specific positioning of residues. On the other hand, 18 cADs (64%) showed activation by the wildtype sequence within 2-fold of the scrambled sequence average, indicating that their activation is primarily determined by aa composition and not by a unique positioning of residues. The wild-type-to-scramble ratio was not correlated with wild-type activation strength (Figure S4A). Instead, cADs with the greatest hydrophobic content had the lowest wild-type-to-scramble ratios (Figure 4B, Pearson’s r = −0.51). Thus, the majority of cADs activate by a composition-driven mechanism that exploits an excess of bulky hydrophobic residues.

**Figure 4.**
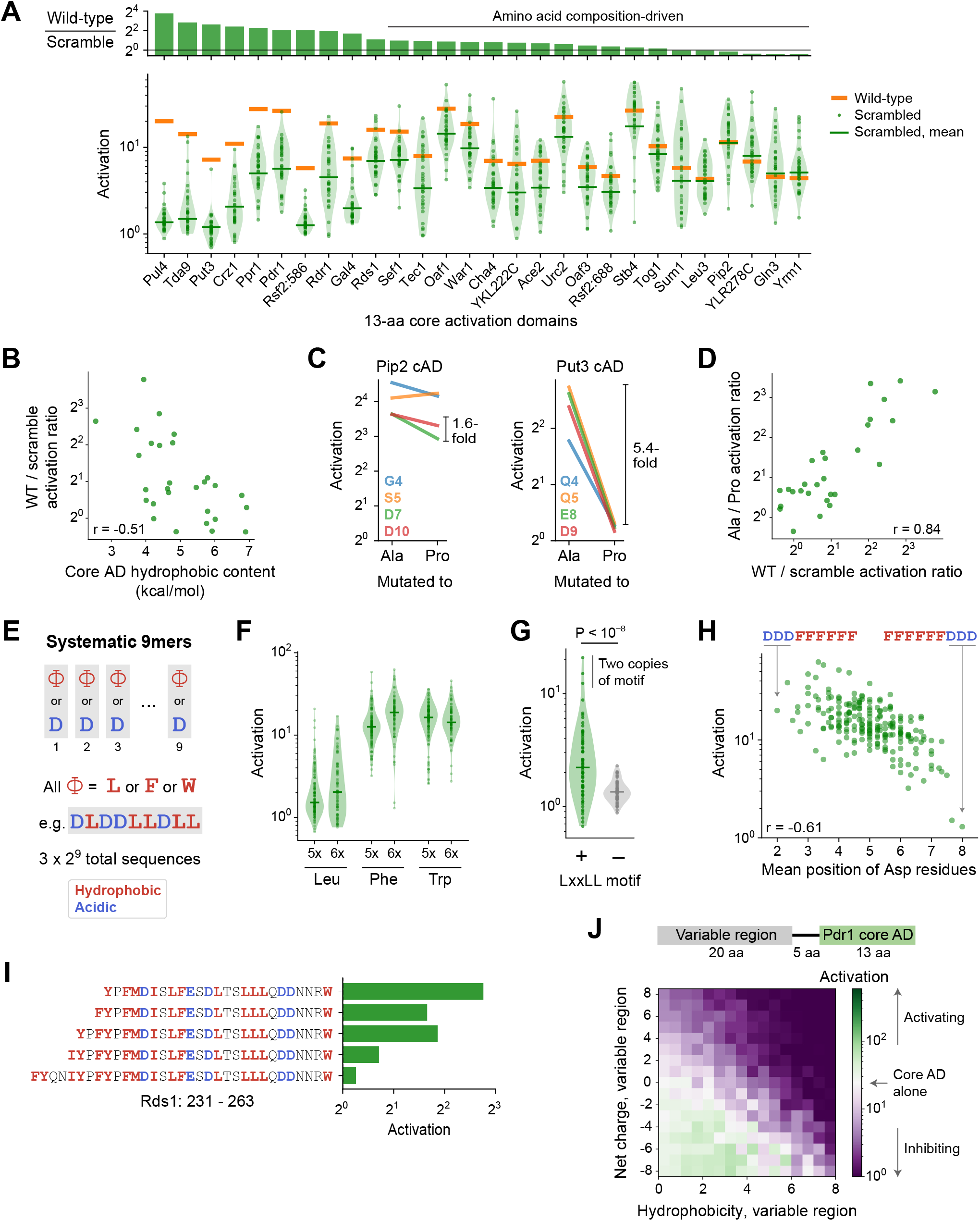
Sequence and structural determinants of activation domains. **(A)** To quantify the importance of aa composition, we measured activation of 33 scrambled sequences for each of the 28 strongest 13-aa core ADs (cAD). (*Top*) Activation of each wild-type cAD divided by the mean activation of its scrambled mutants. Eighteen cADs (64%) showed activation by the wild-type sequence within 2-fold of the scrambled sequence average, indicating that their activation is primarily determined by aa composition and not by a unique positioning of residues. (*Bottom*) Activation of each wild-type cAD (orange bars) and scrambled mutants (green circles), with the mean activation of mutants shown by the green bars. See also Figure S4A. **(B)** cADs with the greatest hydrophobic content had the lowest wild-type-to-scramble ratios. Pearson’s r = −0.51. **(C)** To directly determine whether the 28 strongest 13-aa cADs require helical folding, we individually mutated each non-hydrophobic residue to Ala or Pro and asked whether Pro inhibited activation relative to Ala. Activation of the Pip2 cAD (left) was not disrupted by Pro mutation and showed at most a 1.6-fold drop compared to Ala mutations. In contrast, activation of the Put3 cAD (right) was abolished by all Pro mutations, up to a 5.4-fold drop compared to Ala mutations. Effects of mutations for all 28 cADs are shown in Figure S4B. **(D)** Across cADs, the wild-type-to-scramble ratio (horizontal axis) was correlated with the maximal fold-drop in activation resulting from proline mutation (vertical axis). Pearson’s r = 0.84. **(E)** A simplified system of artificial cADs that uses only the four most activation-promoting residues. Namely, 9-aa sequences (“9mers”) consisting entirely of only two types of amino acids each: Asp and either Leu, Phe, or Trp. This way, all possible sequences (3 x 2^9^ = 1536) could be systematically assayed. **(F)** Activation by all 9mers consisting of five or six Leu, Phe, or Trp residues, grouped by amino acid composition. Activation by all 9mers, grouped by amino acid composition, is shown in Figure S4C. **(G)** Presence of the LxxLL motif is significantly predictive of activation in 9mers with five Leu residues (P < 10^-8^ by Kolmogorov-Smirnov test). The three strongest-activating sequences contain two copies of the motif. This is the only motif strongly associated with activation of these Leu 9mers; effects for all motifs tested are shown in Figure S4D. Similar motifs were also significantly but more weakly associated with Phe 9mers and Trp 9mers (Figure S4E). **(H)** Activation by 9mers with five or six Phe residues was correlated with the average position of their acidic residues (Pearson’s r = −0.61). Most dramatically, DDDFFFFFF activated 20-fold while its reverse sequence FFFFFFDDD activated just 1.3-fold. **(I)** Residues 239-263 alone in Rds1 activated 6.8-fold, but extending the sequence by eight aa, five of them hydrophobic (red), abolished activation. **(J)** (*Top*) To systematically quantify inhibition, we measured the effect on activation when the Pdr1 cAD was placed next to a library of 2177 random 20-aa sequences chosen to span a wide range of net charge and hydrophobic content. (*Bottom*) Sequences are binned by net charge and hydrophobic content of the variable region and mean activation is plotted in color. Activation of the Pdr1 cAD alone is shown in white, with stronger and weaker activation shown in green and purple respectively.

Alpha-helical folding is thought to be important for activation, which is at odds with the prevalence of composition-driven cADs. We directly determined whether the 28 strongest 13-aa cADs require helical folding by measuring the impact of inserting a helix-breaking proline. Within the seven central positions of each 13-aa cAD, we individually mutated each non-hydrophobic residue to alanine or proline and asked whether proline inhibited activation relative to alanine (Figure 4C). Proline mutations inhibited activation more than 3-fold over alanine in nine ADs (32%), including up to an 11-fold drop in activation in the Tda9 cAD. However, proline mutations inhibited activation by less than 2-fold in 16 cADs (58%). These effects were consistent between different positions within the same cAD (Figure S4B). Interestingly, all three cADs that contained a basic residue (Pul4, Tda9, and Rsf2:586) were strongly inhibited by proline, suggesting that a helix is necessary to position their inhibitory positive charge away from the coactivator binding interface. Most notably, the magnitude of proline disruption was tightly correlated with the wild-type-to-scramble ratio (Figure 4D, Pearson’s r = 0.842), showing that constraints on structure and sequence, or a lack thereof, were closely linked. Thus, composition-driven cADs do not need to form a helix and likely sample disordered conformations, while cADs that require their wild-type sequence also require helical folding, employing a structure-driven mechanism.

To understand these two contrasting mechanisms, we studied them in a simplified system of artificial cADs that used only the four most activation-promoting residues. We examined 9-aa sequences (denoted “9mers”) consisting entirely of only two types of amino acids each—Asp and either Leu, Phe, or Trp—so that all possible such sequences (3 x 2^9^ = 1536) could be systematically assayed (Figure 4E). As expected, activation required a balance of hydrophobic and acidic residues (Figure S4C), so we focused our analysis on aa compositions with the strongest median activation: 9mers with five or six hydrophobic residues. Within these, nearly all Phe 9mers and Trp 9mers activated regardless of their sequence (Figure 4F), characteristic of a composition-driven mechanism and consistent with the highly hydrophobic nature of Phe and Trp residues. However, Leu 9mers spanned a wide range of activity with only a minority that were strongly activating, characteristic of structure-driven activation requiring a specific ordering of residues.

To see what allows only certain Leu 9mers to activate, we examined all possible short motifs containing two or three Leu residues. The only motif strongly associated with activation was LxxLL (Figures 4G and S4D; P < 10^-8^ by Kolmogorov-Smirnov test). This motif was strictly required for activation in these 9mers, present in two copies in the three strongest 9mers, found in two wild-type structure-driven cADs (Put3 and Crz1), and previously described in ADs of many nuclear receptors and other TFs (Plevin et al., 2005). This motif also suggests an explanation for why helical folding was necessary for cADs with less hydrophobicity: by placing Leu residues along one face of an alpha helix, the motif efficiently forms a hydrophobic binding surface.

The analogous FxxFF and WxxWW motifs were also significantly but more weakly associated with activation of Phe 9mers and Trp 9mers (Figure S4E). However, their strongest predictor of activation was the average position of acidic residues, with N-terminal skew highly favored (Figure 4H). Most dramatically, DDDFFFFFF activated 20-fold while its reverse sequence FFFFFFDDD activated just 1.3-fold. One role of N-terminal negative charge could be to neutralize the macroscopic dipole that arises from alpha helix backbone hydrogen bonds, stabilizing helical conformations (Rocklin et al., 2017). Even though a helix is not necessary for composition-driven activation, this effect could reduce the entropic cost of binding to coactivators by promoting transient folding.

### Activation domains are strongly inhibited by nearby clusters of hydrophobic residues

Despite the importance of hydrophobic residues for activation, PADDLE also predicted that some TFs contained cADs whose activation was inhibited by nearby clusters of hydrophobic residues. We confirmed this experimentally for Rds1, in which residues 239-263 alone activated 6.8-fold but extending the sequence by eight aa, five of them hydrophobic, abolished activation (Figure 4I). Inhibition by hydrophobic residues also explains the weak activation of some scrambled 53-aa ADs (Figure S4F).

To systematically quantify inhibition, we measured the effect on activation when the Pdr1 cAD was placed next to a library of 2177 random 20-aa sequences chosen to span a wide range of net charge and hydrophobic content (Figure 4J). As expected, negative charge with moderate hydrophobicity bolstered activation. Positive charge without hydrophobicity inhibited activation, but even variable regions with +8 charge (+4 net charge) still activated an average of 4.4-fold. Therefore, the local clustering of negative charges near hydrophobic residues renders the Pdr1 cAD resistant to global changes in net charge, suggesting that interactions between opposite charges within the peptide do not fully prevent binding to coactivators. In contrast, hydrophobic residues at high density inhibited activation completely. This suggests that inhibitory hydrophobic clusters interact with hydrophobic residues in the cAD, completely preventing their interaction with coactivators. Consistent with two independent mechanisms of inhibition, basic and hydrophobic residues together reduced activation additively. In total, these experiments mapped the biochemical and structural properties that drive both activation and inhibition.

### The large majority of activation domains bind Mediator, and its recruitment is a key driver for activation

The surprising ability of simple biochemical features to explain a large and diverse set of ADs suggests that these features may also drive interactions with coactivator complexes. To gain a mechanistic understanding of the AD-coactivator contacts that underlie activation, we mapped the interactions of wild-type and mutant ADs with Mediator. We focused on Med15, the primary subunit of Mediator targeted by ADs in budding yeast, which contains four tandem activator-binding domains (ABDs) (Herbig et al., 2010; Thakur et al., 2008). We isolated the N-terminal portion of Med15 consisting of its four ABDs (hereafter just called “Med15”), confirmed its *in vitro* interaction with Gcn4, and used it for binding experiments (Figure S5A).

To measure binding in high-throughput, we used mRNA display (Takahashi and Roberts, 2009), expressing our library of TF tiles as a pool of protein fragments covalently tagged with their mRNA sequences, and using this pool in Med15 pull-down experiments (Figure 5A). Direct counts of Med15-bound and input protein molecules were obtained by amplifying and sequencing their mRNA/cDNA tags and using unique molecular identifiers to remove PCR duplicates. Finally, the fractional pull-down of each protein fragment was computed by its relative abundance in the Med15-bound sample versus input, normalized to total library concentrations measured by qPCR (Methods). Pull-down measurements were reproducible between replicates (Pearson’s r > 0.93) and consistent between identical protein fragments encoded by distinct but synonymous mRNA sequences (Figures S5B-C). Fragments known to bind Med15 enriched up to 17-fold above random sequence controls and 1000-fold higher than in mock pull-downs with beads alone (Figure S5D, Table S4).

**Figure 5.**
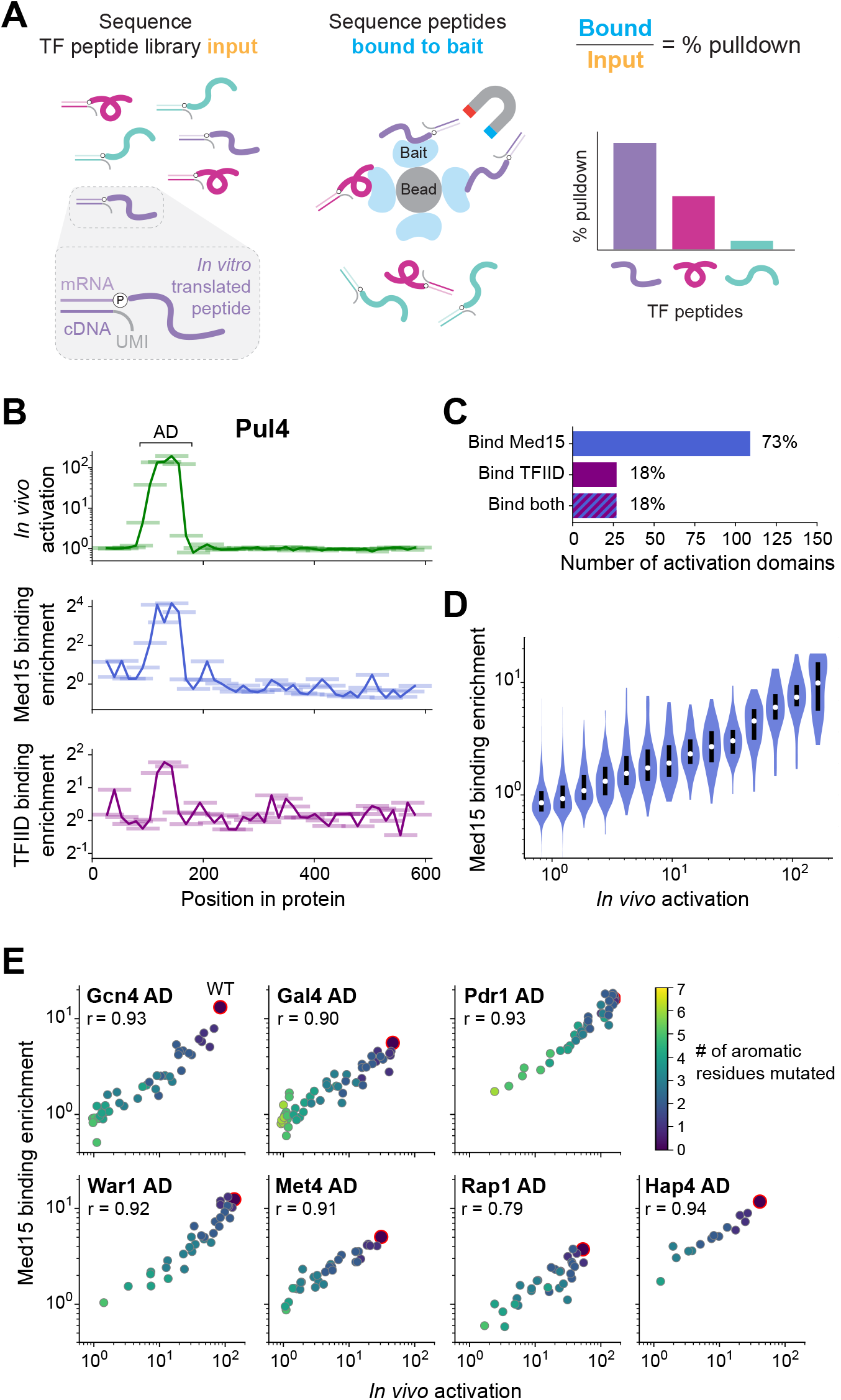
The large majority of activation domains bind Mediator, and its recruitment is a key driver for activation. **(A)** To measure binding in high-throughput, we used mRNA display, expressing our library of TF tiles as a pool of protein fragments covalently tagged with their mRNA sequences (left), and using this pool in pull-down experiments (middle). Direct counts of bound and input protein molecules were obtained by amplifying and sequencing their mRNA/cDNA tags and using unique molecular identifiers (UMIs) to remove PCR duplicates. Finally, the fractional pull-down of each protein fragment was computed by its relative abundance in the bound sample versus input, normalized to total library concentrations measured by qPCR (right). See also Figure S5. **(B)** For all tiles spanning Pul4, *in vivo* activation (green), *in vitro* Med15 binding enrichment (blue), and *in vitro* TFIID subcomplex binding enrichment (purple) are plotted. Tiles, 53 aa long, are shown as horizontal bars with a line traced through their centers for clarity. The Pul4 AD binds both Med15 and TFIID. Binding enrichments of all tiles are in Table S4. **(C)** Number and percentage of ADs that bound Med15, TFIID subcomplex, or both in pull-down experiments. ADs and Med15-binding domains overlapped substantially. All ADs that bound TFIID also bound Med15. See also Figure S5K and Table S4. **(D)** Violin plot of the Med15 binding enrichment of fragments, grouped by their *in vivo* activation. A white dot labels the median of each bin and a black bar marks the 25^th^ to 75^th^ percentile interval. See also Figure S5F. **(E)** We measured Med15 binding of the set of AD mutants in which aromatic residues were systematically removed (Figure 2F). Med15 binding and activation are plotted, with wildtype sequences outlined in red, the number of mutated residues shown in color, and Pearson’s r displayed.

Binding to Med15 was widespread among ADs and coincident with activation activity. In total, 324 tiles bound Med15 significantly more than negative controls (P < 0.001 by Z-test, Figure S5E, Methods). When plotted by protein position, Med15-binding tiles clustered into 153 discrete domains, the vast majority of which were not previously known to bind Mediator (Figure 5B, Table S4). ADs and Med15-binding domains overlapped substantially: 73% of ADs bound Med15, and 71% of Med15-binding domains functioned as ADs (Figure 5C). Moreover, ADs that did not bind Med15 tended to activate weakly (Figure S5F), so it is possible that many bind Med15 at levels below the detection limit of our pull-down assay, which was less sensitive than our activation assay (compare Figures 1C and S5E). Just as with activation, high acidic and hydrophobic content were associated with Med15 binding (Figure S5G). The Gln3 AD that did not require acidic residues also did not bind Med15 substantially (2.1-fold enrichment), suggesting that it activates through other mechanisms.

Most notably, activation strength and Med15 binding of tiles were directly proportional (Figure 5D). To test this relationship further, we measured Med15 binding of the set of AD mutants in which aromatic residues were systematically removed (Figure 2F). In every case, Med15 binding and activation were strongly correlated (Figure 5E), showing that the hydrophobic features that drive activation similarly underlie Med15 binding. Moreover, the ability for binding of a single protein *in vitro* to quantitatively explain activation *in vivo* suggests that Mediator recruitment is a key driver for gene activation and determined largely by AD binding its Med15 subunit.

ADs can bind different coactivators, though the extent of such interaction is unknown. To explore the possibility, we also examined binding of TF tiles to TFIID, a key factor in the transcription of nearly all genes (Warfield et al., 2017). TFIID contains three known AD-binding subunits, Taf4, Taf5, and Taf12, which form a subcomplex with Taf6 and Taf9 (Layer et al., 2010; Wright et al., 2006). We isolated this TFIID subcomplex and confirmed its interaction with the VP16 AD (Figure S5H). Pull-downs of the TF tiling library with the TFIID subcomplex enriched fragments up to 6.7-fold above random sequence controls (Figures S5I-J, Table S4). In total, 27 ADs (18%) across 26 TFs bound the TFIID subcomplex (P < 0.001), greatly expanding its repertoire of known interactions (Figures 5B-C). However, all ADs that bound TFIID also bound Med15, and despite the identical pull-down conditions, ADs bound more weakly to TFIID than to Med15 (Figure S5K). While more TFIID interactions may be discoverable if assay sensitivity can be further optimized, these results suggest that TFIID has broadly lower affinity for ADs and provide no evidence for its specific targeting as a coactivator. Since TFIID binding by ADs is redundant and less frequent, its role in TF-directed activation is at most secondary to that of Mediator.

### Med15 uses a shape-agnostic, fuzzy interface to bind diverse activation domain sequences

What structural features of Med15 enable its promiscuous yet functional interaction with diverse AD sequences? In the best studied example, the Gcn4 AD interacts with hydrophobic patches and basic residues on multiple Med15 activator-binding domains (ABDs) in a large number of binding poses to form a “fuzzy” complex (Brzovic et al., 2011; Tuttle et al., 2018). We addressed the question by using FlexPepDock, a peptide docking algorithm from the Rosetta suite, to systematically build structural models of ABD-AD interactions, made computationally tractable by our identification of short cADs (Leaver-Fay et al., 2011; Raveh et al., 2011). Our modeling focused on two ABDs with known structures: the KIX domain, which has homology to human Med15 and the p300/CBP coactivator family, and ABD1 (Figure 6A) (Brzovic et al., 2011; Thakur et al., 2008). To sample diverse sequences, aa composition, and secondary structure, we modeled interactions of the 28 13-aa cADs described above (Figures 4A-D) with the KIX domain and ABD1. For each interaction, 50,000 candidate structural models were sampled and ranked by the Rosetta energy score, and the 10 best-scoring models from each interaction were used in subsequent analyses (Table S5).

**Figure 6.**
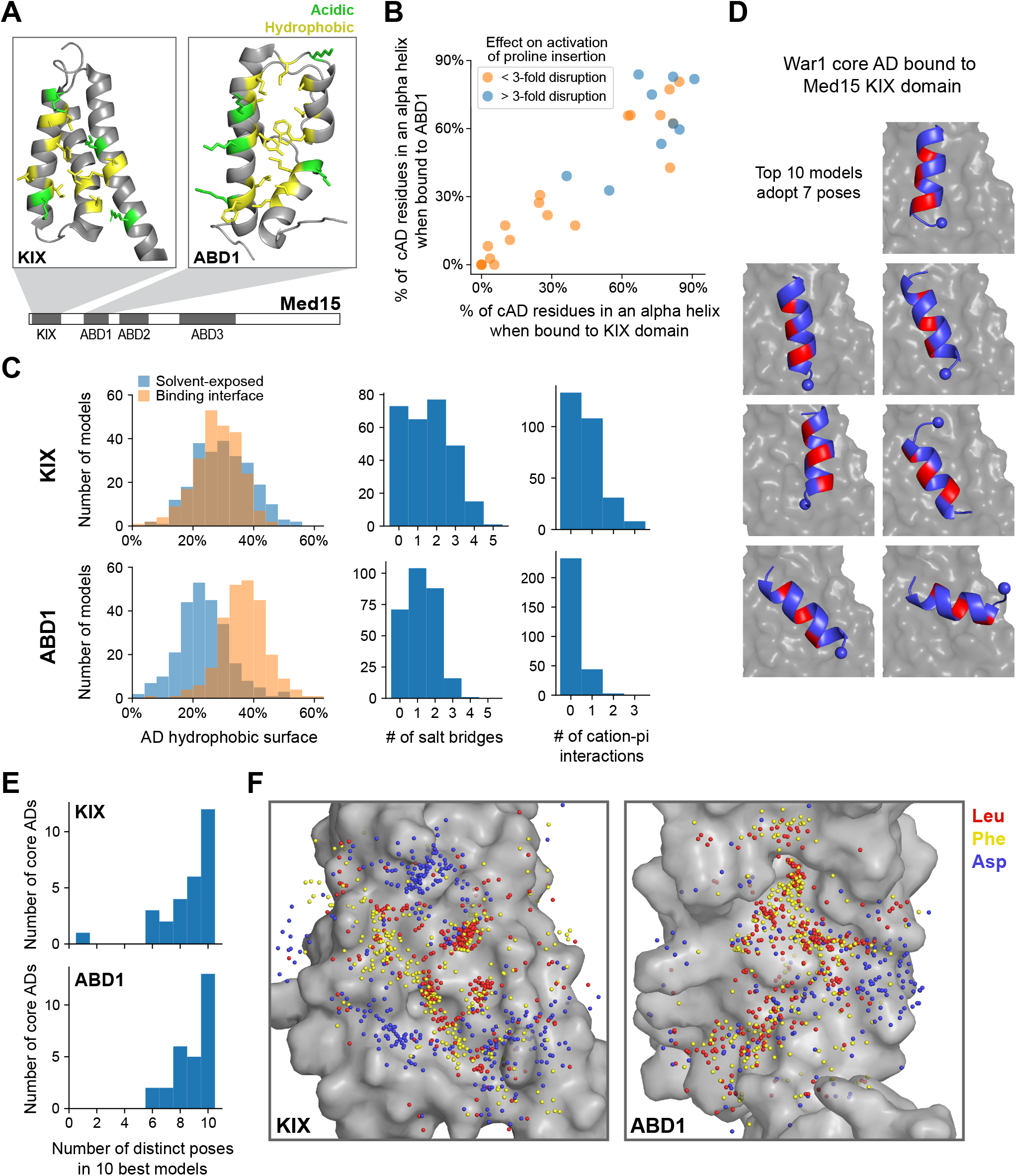
Med15 uses a shape-agnostic, fuzzy interface to bind diverse activation domain sequences. **(A)** We used a Rosetta peptide docking algorithm (Raveh et al., 2011) to build structural models of the 28 13-aa core ADs (cADs) described above (Figures 4A-D) interacting with two activator-binding domains (ABDs) of Med15, the KIX domain and ABD1. Structures of these domains are shown, with the hydrophobic (yellow) and acidic (green) residues that form the AD-binding surfaces displayed (Thakur et al., 2008; Herbig et al., 2010). For each interaction, 50,000 candidate structural models were sampled and ranked by the Rosetta energy score, and the 10 best-scoring models from each interaction were used in subsequent analyses (Table S5). See also Figure S6A. **(B)** The percentage of cAD residues that form an alpha helix in the 10 best-scoring models when bound to the KIX domain or ABD1. cADs that were more than 3-fold disrupted by mutations to proline (Figure 4C) are colored blue. **(C)** Histograms summarizing the structural features of the 10 best-scoring models of all cADs bound to the KIX domain (top) and ABD1 (bottom). *(Left)* Fraction of the total hydrophobic surface of cAD residues that is solvent exposed (blue) or at the binding interface (orange). See also Figure S6B. *(Right)* Number of salt bridges or cation-pi interactions formed between cADs and the KIX domain or ABD1. **(D)** The 10 best-scoring models of the War1 cAD (blue) bound the KIX domain (grey surface) with its helix axis at different orientations and with different helical faces presented towards the interaction surface, in seven distinct poses in total. For orientation, the cAD N-terminus is shown as a sphere and one helical face is colored red. See also Figure S6C. **(E)** Distinct binding poses were defined by clustering similar structures based on cAD backbone root mean square distance, and the number of poses seen in the 10 best-scoring models was counted for cAD interactions with the KIX domain (top) and ABD1 (bottom). See also Figures S6C-E. **(F)** Sidechain locations of all Leu (red), Phe (yellow), and Asp (blue) residues of all cADs interacting with the KIX and ABD1 surfaces (grey) are marked by dots. Leu and Phe distributions were similar to each other: there was no binding pocket that selectively preferred one residue over the other, despite large differences in their size and shape.

We validated our structural modeling by comparison to experimental data in two ways. First, KIX domain residues previously shown by NMR to be important for interacting with the Pdr1 AD were recapitulated by the best-scoring model of this interaction (Figure S6A) (Thakur et al., 2008). Second, the degree of alpha helix formation across the 28 cADs was consistent with the effect of proline insertion on *in vivo* activation (Figure 6B). As expected, cADs that were inhibited by proline predominantly bound both domains as an alpha helix, and conversely cADs that bound both domains in disordered conformations were minimally affected by proline insertion.

A unifying feature of the interactions was the prominent role of hydrophobic contacts at the protein interface. Hydrophobicity-driven interactions between proteins often employ high shape complementarity to maximize interface area and minimize solvent-exposed area. However, the flat surface of the KIX domain was ineffective in burying hydrophobic cAD residues, which on average had as much surface area exposed to solvent as area contacting the KIX domain (Figure 6C, top). For example, the best-scoring Pdr1 AD model used Ile and Leu residues along one helical face to contact the KIX domain, but this left Trp and Tyr residues on the opposite face exposed to solvent (Figure S6B). The poor hydrophobic shape complementarity of the KIX domain was frequently mitigated by multiple salt bridges and cation-pi interactions (Figure 6C). ABD1, which formed fewer ionic interactions, engaged cADs through a larger surface formed by its contoured hydrophobic ridge (Figure 6C, bottom). Nevertheless, hydrophobic residues of most cADs were incompletely buried, with those residues showing greater than 20% of area exposed to solvent in over half of all structures. Thus, despite the central role of hydrophobic AD residues in activation and Med15 binding, hydrophobic shape complementarity is not an important feature of ABD-cAD interactions.

Consistent with weak constraints on the shape of the binding interface, all cADs bound in diverse poses. For example, the 10 best-scoring models of the War1 cAD bound the KIX domain with its helix axis at different orientations and with different helical faces presented towards the interaction surface (Figure 6D). To quantify this diversity, we defined the unique binding poses of each ABD-cAD interaction by clustering similar structures based on cAD backbone root mean square distance (2 Å distance cutoff), and then counted the number of poses adopted by the 10 best-scoring models. The 10 best-scoring models of nearly every cAD, when interacting with either domain, adopted 6 or more distinct poses (Figure 6E). To ensure that this did not result from insufficient sampling, we generated ten-fold more candidate models (500,000 total) for six cADs binding the KIX domain. For all six cADs, the 10 best-scoring models still adopted 7 or more distinct poses, despite spanning a narrower range of Rosetta scores (Figures S6C-D). Disordered cADs, lacking helical content, adopted especially many distinct poses; also, poses were frequently sampled only once, suggesting that disordered cADs inhabit an extremely large conformational space that may reduce the entropic cost of binding (Figures S6E-F). Altogether, these structures point to a shape-agnostic nature of Med15 ABD interaction surfaces and support the idea of a multi-pose, fuzzy mode of interaction.

To explore the consequence of fuzzy binding for amino acid sequence constraints, we took Leu, Phe, and Asp residues from models of all cADs and plotted their binding positions on the KIX and ABD1 interfaces (Figure 6F). As expected, the hydrophobic Leu and Phe residues occupied regions distinct from the negatively charged Asp residues. However, the Leu and Phe distributions were similar to each other: there was no binding pocket that selectively preferred one residue over the other, despite large differences in their size and shape. The diversity of AD sequences may thus be understood in terms of the fuzzy nature of the AD-ABD interaction, which only loosely constrains the characteristics of hydrophobic AD residues.

### The high valence of TF-Mediator interactions enables tunable, long-lived, yet dynamic binding

Pull-down experiments with individual ABDs showed no appreciable binding to any of nine ADs (Figure S7A), consistent with the previous suggestion that multiple ABDs are required for strong binding to Med15 (Brzovic et al., 2011; Tuttle et al., 2018). Multiple interaction sites were also common within ADs: PADDLE identified 42 ADs (28%) that contained two or more non-overlapping cADs (Figure 7A, Table S3, Methods). We took 47 pairs of adjacent cADs and measured activation by the two cADs individually or in tandem. For 40 of the pairs, the tandem cADs activated more strongly than expected from the combined activation of the individual cADs (4.0-fold median increase, Figure 7B). Thus, an increased valence of interaction sites in both Med15 and ADs drives stronger binding and activation.

**Figure 7.**
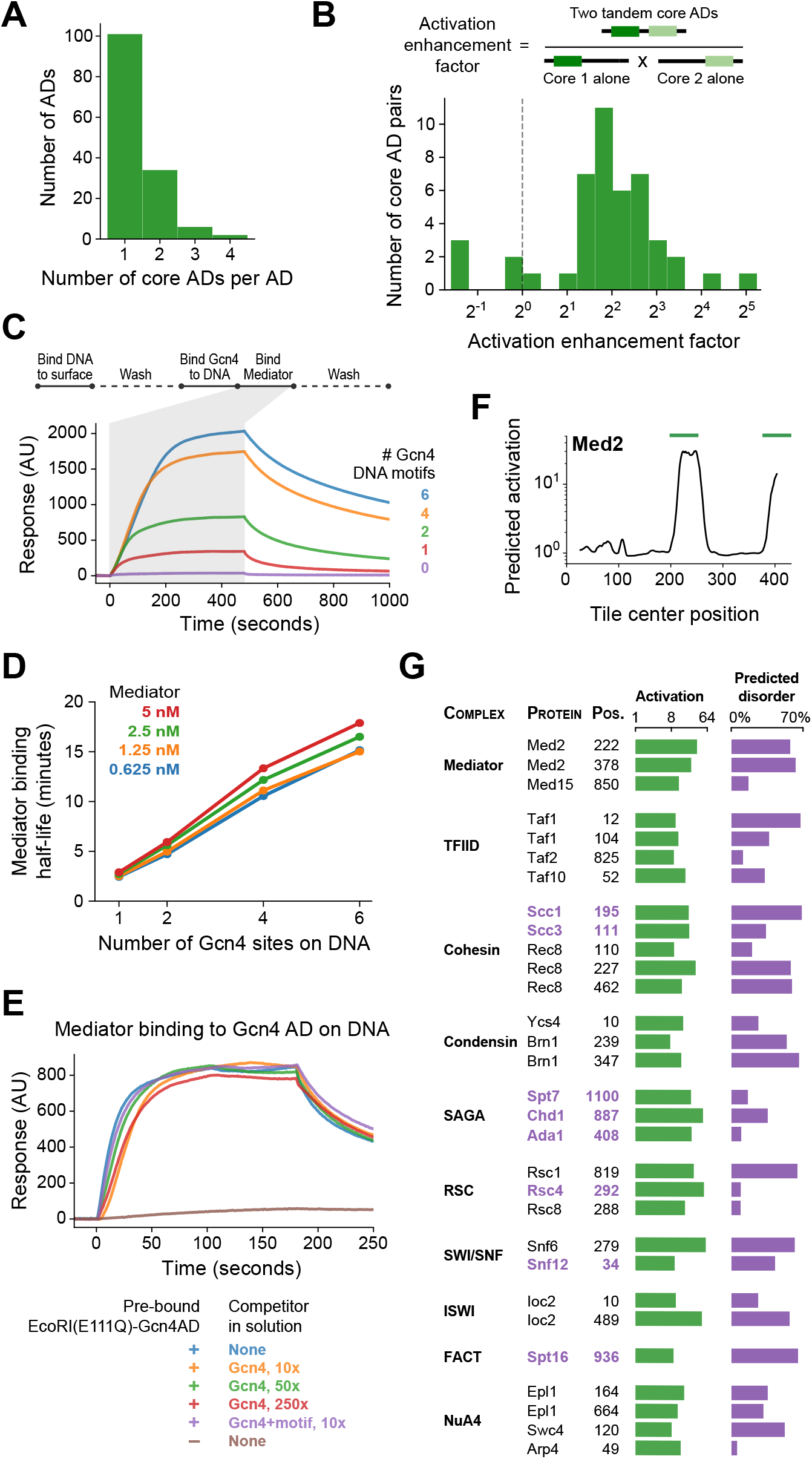
Functional consequences of high-valence coactivator interactions. **(A)** PADDLE identified 42 ADs (28%) that contained two or more non-overlapping cADs. **(B)** We took 47 pairs of adjacent cADs and measured their activation enhancement factors—the activation of both cADs in tandem divided by the product of activation by each cAD individually. For 40 pairs, activation was enhanced when cADs were in tandem, with a median enhancement factor of 4.0-fold. **(C)** Kinetics of Mediator complex recruitment to DNA by Gcn4. (*Top*) We coupled DNA containing a Gcn4 motif to a NeutrAvidin-coated surface, added Gcn4, and measured real-time binding of Mediator by surface plasmon resonance. The Mediator binding step also included 7.5 nM Gcn4 to maintain DNA-bound Gcn4. (*Bottom*) Real-time binding of 2.5 nM Mediator to Gcn4-DNA complexes (0 to 480 seconds) and subsequent dissociation (480 seconds onwards). DNA templates contained 0, 1, 2, 4, or 6 copies of the Gcn4 motif. AU, arbitrary units. **(D)** The interaction half-life of Mediator with Gcn4-DNA complexes was proportional to the number of Gcn4 motifs and independent of the concentration of Mediator used in the binding step. See also Figure S7B-C. **(E)** Gcn4 competition assay. We purified a fusion of the Gcn4 AD to nuclease deficient EcoRI(E111Q), which resides on DNA for several hours, bound it to DNA containing a single EcoRI site, and measured Mediator recruitment in the presence or absence of excess Gcn4 competitor. Mediator was at 5nM in all six conditions. AU, arbitrary units. See also Figure S7D. **(F)** PADDLE predictions on 53-aa tiles spanning yeast Med2 identified two ADs, marked with green bars, which were tested experimentally. An activation screen of nuclear proteins identified ADs in all major coactivator and chromatin modifying complexes. Protein complex, protein name, and start position of the 53-aa AD is labeled, and experimentally-measured activation (green) and fraction of residues predicted to be disordered (purple; in D2P2) is shown. AD that are predominantly disordered or unresolved in PDB structures are displayed in purple text. See also Figure S7E and Table S1.

At larger scales, the number of interaction sites is further multiplied because some TFs contain multiple ADs, many TFs bind DNA as dimers, and many genes have several TF binding sites (Hahn and Young, 2011; Spitz and Furlong, 2012). To understand the advantages conferred by the high valence of TF-Mediator interactions, we studied the kinetics *in vitro* of the initial step of activation, TF-driven recruitment of Mediator to DNA. We performed these experiments with Gcn4, which has an extended AD consisting of three cADs, and which dimerizes through its DBD. We isolated budding yeast Mediator complex, coupled DNA containing a Gcn4 motif to a NeutrAvidin-coated surface, added Gcn4, and measured real-time binding of Mediator by surface plasmon resonance (Figure 7C). Mediator bound to the Gcn4-DNA complex with an apparent dissociation constant of K_D_ = 14 nM ± 2 nM and an interaction half-life of 2.7 ± 0.2 minutes (Figure S7B). We repeated the experiment with DNA containing 2, 4, or 6 copies of the Gcn4 motif. Mediator bound the resulting Gcn4-DNA complexes with identical on-rates. Mediator was also recruited more efficiently, with more Mediator bound per mole of Gcn4 than on templates with only one motif (Figure S7C). The interaction half-life, however, increased with additional copies of the Gcn4 motif—up to 16 minutes when 6 motifs were present (Figures 7C-D). If all copies bound Mediator independently, then the half-life should be the same, regardless of the number of copies. The presence of additional, neighboring ADs retards dissociation by enabling recapture of Mediator released from one AD through binding to another. A slower dissociation rate corresponds to an increase in apparent affinity constant.

Conversely, we found that the multiplicity of ABDs in Mediator facilitated binding of Gcn4 ADs. When the surface plasmon resonance experiment was repeated with large excess Gcn4 in solution, there was almost no change in the measured on-rate. Even a 160-fold excess of Gcn4 had only minimal effect on the rate of Mediator binding and final level achieved (Figure S7D). To maintain stable TF-DNA complexes without TFs in solution, we repeated these experiments using a fusion of the Gcn4 AD to nuclease deficient EcoRI(E111Q), which resides on DNA for many hours and also forms homodimers (Wright et al., 1989). Still, Mediator with no Gcn4 competitor or with 250-fold excess of Gcn4 competitor bound similarly (Figure 7E). Excess competitor Gcn4-DNA complexes also failed to slow Mediator binding to the surface. Evidently, binding of Gcn4 to Mediator does not appreciably block binding of other Gcn4 molecules. This behavior may be explained by the occurrence of multiple Gcn4-binding sites on Mediator and weak interaction of a cAD with any one site. Rapid association-dissociation equilibrium of a cAD allows a second Gcn4 molecule to interact with Gcn4-bound Mediator, increasing the frequency of exchange of one Gcn4 molecule by another. Such facilitated exchange provides a rationale for both the fuzzy nature of the AD-ABD interaction and the multiplicity of ABDs.

### Activation domains are present in all major coactivator complexes

PADDLE predictions and previous experiments (Liu and Myers, 2015) identified two strong ADs in the Med2 subunit of Mediator, which is adjacent to Med15 in the Mediator complex, suggesting a broader role for ADs beyond TF proteins (Figure 7F). We used PADDLE to search for ADs across all nuclear-localized non-TF proteins in yeast and selected 1485 fragments that showed potential for activation (Methods). Upon pooled *in vivo* testing, we found 290 fragments across 229 proteins that activated significantly (P < 0.001 by Z-test, Table S1). To see how often PADDLE fails to identify acidic ADs, we also included 291 control fragments that had extremely high acidic and hydrophobic content but were predicted not to activate. No control fragments activated (Figure S7E), showing that PADDLE likely identified all acidic ADs that exist and again demonstrating that AD function cannot be predicted from amino acid composition alone.

ADs were remarkably widespread and were found in proteins associated with diverse functions including transcriptional regulation, RNA splicing, DNA repair, ribosome biogenesis, and protein degradation. We focused on the ADs in transcription-associated proteins. Fourteen ADs were found in proteins that bind DNA and regulate pathway-specific genes; these were not included in the original tiling library because they bind DNA only through a TF partner. The two predicted Med2 ADs were also confirmed to activate, 36-fold and 25-fold in our assay.

Notably, ADs were identified in all major coactivator and chromatin modifying complexes, including Mediator, TFIID, cohesin, condensin, SAGA, RSC, SWI/SNF, ISWI, NuA4, and FACT (Figure 7G). These complexes contained 2.9-fold more activating fragments than expected by chance (P < 10^-12^ compared to randomly shuffled controls), suggesting a functional role. Domains that activate on their own could be inactive in their natural context if buried within a protein or protein complex. To investigate this possibility, we crossreferenced coactivator ADs with all available structures in the Protein Data Bank (PDB) and with the Database of Disordered Protein Predictions (D2P2) (Berman, 2000; Oates et al., 2012). In fact, 23 ADs were predicted in D2P2 to be highly disordered (more than 30%) or were predominantly disordered or unresolved in PDB structures (Figure 7G), and so may be exposed and active *in vivo.* For example, cohesin, which binds Mediator and forms enhancer-promoter contacts through loop extrusion (Davidson et al., 2019; Kagey et al., 2010; Kim et al., 2019; Sanborn et al., 2015), contained strong and likely-disordered ADs in its subunits Scc1 (RAD21 homolog), Scc3 (STAG1/2 homolog), and Smc3, as well as multiple ADs on the meiotic-specific subunit Rec8. These cohesin ADs, as well as ADs within other coactivators, may play a role in gene regulation by driving interactions with Mediator.

## DISCUSSION

Although activation domains (ADs) have been studied for over three decades, their locations in transcription factors (TFs) and mechanisms of activation have remained enigmatic. Combining quantitative, high-throughput experiments and computational modeling, we determined the biophysical principles underlying activation by acidic ADs. We trained a neural network to predict ADs in yeast and humans, which led to the discovery of new ADs and activation mechanisms. While an amphipathic helix can enhance activation, most ADs simply activate through an abundance of clustered acidic and bulky hydrophobic residues. ADs bind the hydrophobic and basic surface of Med15 activator-binding domains (ABDs) through a fuzzy interface. The low shape complementarity of this interaction only weakly constrains AD sequences, explaining their high diversity. We show that the dynamic nature of transcriptional signaling arises from the fuzzy and multivalent nature of TF-Mediator interactions. Finally, an expanded role for ADs is suggested by our finding that ADs are present in all major coactivator complexes.

### Quantitative, high-throughput measurements of activation enabled prediction of new ADs and revealed the principles underlying acidic activation

We obtained high-throughput measurements of activation strength that were both quantitative and precise. This enabled us to detect—and predict with PADDLE—how activation strength depends on amino acid sequence, amino acid composition, and valence. Quantitation was achieved by inducing artificial TF (aTF) binding for a brief defined period so that GFP levels were representative of transcriptional activation (McIsaac et al., 2013), and by partitioning the range of GFP signal into eight levels for sorting. Others have measured activation only as an on-off binary system or screened for non-quantitative measures of activation, such as cell survival or pull-down of a cell surface marker (Arnold et al., 2018; Ravarani et al., 2018; Tycko et al., 2020). We achieved high precision by restricting measurements to cells with a defined level of aTF expression. Controlling for this frequently-overlooked confounding variable was critical because activation was strongly dependent on aTF abundance, especially at low levels of expression (Figure S1E). Another approach is to account for TF expression by sorting based on the GFP reporter signal divided by aTF expression (Staller et al., 2018); this introduces substantial variability, because activation has a non-linear relationship with aTF expression (Figure S1E), and because this ratio can be systematically biased by features that affect aTF expression (Figure S1F). This may be the reason why it has been concluded that acidic residues are not necessary for Gcn4 activation, whereas using a similar reporter system, we found definitive evidence to the contrary.

By this approach, we found that all ADs in *S. cerevisiae* employ a shared acidic and hydrophobic basis and trained a neural network termed PADDLE to predict sequences that activate on this basis. PADDLE predicted ADs in human TFs with 92% accuracy, indicative of broad conservation of the acidic activation mechanism. PADDLE identified hundreds of new ADs in human TFs and virus proteins. PADDLE could further be used to interpret the functional impact of cancer-associated and naturally-occurring mutations in human ADs.

During our development of PADDLE, a predictor of yeast activation named “ADpred” was published (Erijman et al., 2020). ADpred, a shallow neural network trained on random protein sequences, achieved a prediction accuracy of 93% on random sequences. However, its accuracy on wild-type sequences was much lower: only 33% of ADs predicted by ADpred activated in our experiments, and 29% of the ADs we identified were not predicted by ADpred, only 3.1-fold and 2.3-fold better than random predictions, respectively (alternatively, tile-wise AUPRC of 0.294; Figures S3I-J, Methods). This discrepancy in accuracy may be because the random sequences on which ADpred was trained do not sufficiently represent the vastly larger space of actual protein sequences; for example, random sequences rarely form significant secondary structure. This failure to generalize highlights the importance of matching neural network training data to the prediction task and underscores the advantage of experiments on wild-type and related mutant sequences.

PADDLE predictions were also crucial for generating, refining, and testing hypotheses for activation mechanisms. For example, we unexpectedly discovered that hydrophobic residues could inhibit activation by noticing that subdomains of certain non-activating regions were predicted to activate on their own. More generally, predictions on arbitrary subsequences of TFs identified the core domains responsible for activation at single-amino acid resolution, focusing subsequent experiments on these domains. By combining a high-throughput assay to measure and a machine learning algorithm to predict activity of protein sequences, we have demonstrated a design-build-test-learn cycle that could be applied to accelerate discovery of protein function in other areas of research.

This approach yielded the most detailed view to date of the principles governing acidic AD sequence, which we summarize as follows. Activation arises from an abundance of acidic and bulky hydrophobic residues, especially Asp, Trp, Phe, and Leu. The most potent activators cluster these residues densely, forming short core ADs (cADs). In most ADs, and especially with high hydrophobicity and abundant Phe and Trp residues, this clustering is sufficient to activate in a composition-driven manner, regardless of sequence or secondary structure. ADs with lower hydrophobic content, particularly those with many Leu residues, instead must position their hydrophobic residues along one face of an alpha helix to activate. Overall, the loose constraints on sequence explain the remarkable diversity and lack of evolutionary conservation of ADs.

The unusual plasticity in AD sequence has several advantages. It facilitates spontaneous evolution of new ADs, since an appreciable proportion of even random sequences can activate. This flexibility would allow organisms to develop new transcriptional circuits responsive to their unique needs. Furthermore, in contrast to classical interactions in which structure and function are dependent on a few key residues, activation strength can be adjusted gradually by mutation. Indeed, the observation that nearly all core ADs could increase their strength with simple mutations demonstrates that proper regulation requires precisely adjusted rather than maximized activation. Similarly, AD sequences can preserve their activity while evolving other features, such as in the Gal4 C-terminal AD, whose sequence is specifically bound and inhibited by Gal80 in the absence of galactose (Johnston et al., 1987; Ma and Ptashne, 1987).

### ADs bind Mediator using a shape-agnostic fuzzy interaction, which explains the heterogeneity of AD sequences and enables facilitated exchange of bound molecules

With the use of mRNA display, we found that 73% of ADs bound the Med15 subunit of Mediator and that binding strength was strongly predictive of activation for thousands of wild-type and mutant protein sequences. In contrast, a five-protein TFIID subcomplex bound only 18% of ADs, all of which also bound Mediator. We conclude that Mediator recruitment by TFs is a key driver for gene activation, and Mediator recruitment is largely determined by affinity of ADs for the Med15 subunit. These findings are in keeping with reports that Mediator recruitment occurs independently of other coactivators and can stimulate the recruitment of other coactivators (Ansari and Morse, 2013; Ansari et al., 2014; Govind et al., 2005).

How is Med15, through its four activator-binding domains (ABDs), able to interact with such a large diversity of AD sequences? Despite the importance of hydrophobic interactions, our structural modeling showed that neither the Med15 KIX domain nor ABD1 provided enough shape complementarity to bury the hydrophobic residues of cADs. Consistent with this, most cADs did not need to form an alpha helix to activate. The lack of shape constraint has two important consequences. First, all modeled cADs bound each ABD in many distinct, equally-favored poses. We therefore suggest that fuzzy binding to Med15, previously demonstrated for the Gcn4 AD, is employed by all ADs. Second, neither ABD showed favored binding positions of Leu versus Phe residues of the cAD, suggesting that the shapes of those residues were unimportant. Thus, the loose constraints on binding shape explain the loose constraints on AD sequence. Instead, each ABD can be approximated as a hydrophobic surface, which engages hydrophobic AD residues through simple hydrophobic forces, flanked by basic residues, which form salt bridges and cation-pi interactions with acidic and aromatic AD residues. Clusters of hydrophobic residues adjacent to ADs inhibit activation, presumably because they compete for interaction with hydrophobic AD residues.

Many experiments have shown that transcriptional initiation involves a cycle in which (1) TFs recruit Mediator to enhancers or upstream activating sites, (2) Mediator contacts the promoter and orchestrates a series of events starting with chromatin rearrangement, followed by pre-initiation complex formation and finally RNA polymerase II recruitment, and (3) Mediator facilitates phosphorylation of the polymerase C-terminal domain, which releases polymerase into transcriptional elongation and triggers dissociation of Mediator (Jeronimo and Robert, 2014; Knoll et al., 2018; Robinson et al., 2016; Whyte et al., 2013; Wong et al., 2014). Mediator release is apparently necessary because stably recruiting Mediator by fusing individual subunits to a DNA-binding domain failed to activate in nearly all subunits tested (Wang et al., 2010). The cycling of Mediator complexes increases the responsiveness of transcriptional regulation in at least two ways. First, it requires certain steps to be repeated in order to continue initiating transcription, preventing the system from locking into an activated state.

Second, it frees Mediator complexes to participate in regulation of other genes, which all depend upon and compete for a relatively limited supply of Mediator (Flanagan et al., 1991; Gill and Ptashne, 1988). Thus, TFs must recruit Mediator in a manner that is specific and high-affinity but still dynamic.

Our kinetic experiments showed that Mediator binds Gcn4-DNA complexes with high affinity but that excess Gcn4 does not impede this interaction, showing that interacting Mediator and Gcn4 molecules can exchange rapidly. If a bimolecular interaction occurs through a single site, a competing molecule cannot bind until the first molecule dissociates which, in a case of high affinity interaction, is a slow process. Our observations are indicative of facilitated exchange, arising from the multiple sites and fuzzy nature of the Gcn4-Mediator interaction. Fuzzy interactions allow for rapid association and dissociation, because a high proportion of random encounters lead to weak but productive binding (Ferreira et al., 2005; Sugase et al., 2007; Tompa and Fuxreiter, 2008). One among multiple binding sites will frequently be available for a second molecule, which can invade and facilitate release of the first. Similarly, accelerated dissociation or exchange of the *E. coli* sequence-flexible DNA-binding protein Fis in the presence of competitor proteins was proposed to occur by transitioning through a destabilizing Fis-DNA-Fis ternary complex (Graham et al., 2011; Kamar et al., 2017). These advantageous kinetics provide a rationale for the prevalence of multivalent fuzzy interactions among transcription proteins.

### Coactivator complexes contain ADs that may drive cross-interaction

ADs have primarily been characterized in TFs where they serve to recruit coactivators such as Mediator, TFIID, SAGA, SWI/SNF, and NuA4 (Hahn and Young, 2011; Mitchell and Tjian, 1989). Our finding that functional ADs are also present in all coactivator complexes suggests that ADs have broader roles. First, coactivator ADs could interact with ABDs within the same complex, limiting their weak or non-specific binding to TFs. These auto-inhibited coactivators could still bind to ADs clustered on DNA through facilitated exchange. For example, we found that two cADs activated more efficiently and two DNA-bound Gcn4 dimers recruited Mediator more efficiently in tandem than individually. Second, coactivator ADs could amplify activation by mediating interactions between coactivators (Ansari and Morse, 2013; Liu and Myers, 2015). For example, upon Mediator binding to TFs, the two ADs on Med2 would be liberated from autoinhibition and could recruit other coactivators. Alternatively, Mediator and other coactivators could cluster in the nucleoplasm through AD-ABD cross interactions and bind together to promoters. Either mode of interaction would explain why recruitment of the Med2/Med3/Med15 subcomplex suffices to recruit SWI/SNF to the *CHA1* promoter and suffices for activation of the *ARG1* gene, while deletion of both Med2 ADs impairs transcription of galactose-inducible genes (Ansari et al., 2014; Liu and Myers, 2015; Zhang et al., 2004).

Phase separation of Mediator and TFs has recently been demonstrated *in vitro* and is proposed to underlie enhancer-promoter clustering and transcriptional activation *in vivo* (Boija et al., 2018; Chong et al., 2018; Sabari et al., 2018; Shrinivas et al., 2019). Phase separation may occur when a protein makes dynamic self-interactions through two or more binding domains, (Banani et al., 2016). Interactions between the four Med15 ABDs and the two Med2 ADs make possible phase separation of the Mediator tail subcomplex; other coactivators, with both ADs and ABDs, might phase separate as well. The functional role of such phase separation, if it occurs, is unclear. A parsimonious interpretation suggests that phase separation of coactivators and TFs is simply a macroscopic consequence of the high valence and dynamism of coactivator-TF interactions, features which may serve an altogether different purpose.

### Solving the enigma of AD structure and function

ADs have long been enigmatic, due to the apparent mismatch of their structure and function: AD sequences are abundant among random polypeptides, and yet ADs bind their targets with high specificity; AD peptides are apparently disordered, and yet they bind with high affinity. We now understand that these structural features of ADs are ideally suited to their functions—they render AD-target interaction dynamic, through rapid yet weak binding to single sites and strong binding yet rapid displacement from multiple sites. Dynamic AD-target interaction may serve various purposes, such as a rapid response to changing conditions, and the recruitment of multiple AD-bearing proteins to a single target. For example, Mediator bound to RNA polymerase II at a promoter might interact transiently with TFIID, SAGA, SWI/SNF complex and other proteins during transcription. This mechanism may be employed with other unstructured sequences and high valence targets in other cellular processes.

## ACKNOWLEDGEMENTS

We thank Ramon Lorenzo Labitigan, Peter Geiduschek, and members of the Kornberg lab for critical feedback and discussions; Ramon Lorenzo Labitigan for assistance with the manuscript and figures; Ralph Davis and Shigeki Nagai for providing purified Mediator; Barbara Davis for assistance with experiments; Namita Mitra and Zane Colaric for sequencing; Ricardo Zermeno and the Stanford Shared FACS Facility for cell sorting; the Weis lab for sharing tissue culture facilities; Rich Roberts and Terry Takahashi for advice on mRNA display; the Stanford Research Computing Center for providing computational resources and support; and Shantao Li, Haiwen Gui, Sarah Gurev, Avanti Shrikumar, and Christopher Yeh for help with machine learning. This work was supported by NIH grants R01-DK121366 and R01-AI021144 (to R.D.K.); NIH grant R01-GM127359 and the U.S. Department of Energy (DoE), Office of Science, Office of Advanced Scientific Computing Research, Scientific Discovery through Advanced Computing (SciDAC) program (to R.O.D.); an NSF Physics Frontiers Center Award (PHY1427654), the Welch Foundation (Q-1866), a USDA Agriculture and Food Research Initiative Grant (2017-05741), an NIH 4D Nucleome Grant (U01HL130010), and an NIH Encyclopedia of DNA Elements Mapping Center Award (UM1HG009375) (to E.L.A.). A.L.S. was supported by the U.S. Department of Defense through the National Defense Science & Engineering Graduate (NDSEG) Fellowship Program. R.J.L.T. is supported by the DoE, Office of Science, Office of Workforce Development for Teachers and Scientists, Office of Science Graduate Student Research (SCGSR) program under contract no. DE-SC0014664. C.V.H. was supported by the Stanford Bio-X Undergraduate Summer Research Program. J.T.F. is supported by NIH grant F32-GM126704. Stanford may seek to commercialize aspects of this work, and related applications for intellectual property have been filed.

## Author contributions

A. L.S. conceptualized the project, developed the methodology, and analyzed data. A.L.S. and C.V.H. performed experiments. A.L.S., B. T.Y., and C.V.H. performed bioinformatic analyses. A.L.S. and B.T.Y. developed machine learning models. A.L.S., J.T.F., and C.V.H. purified proteins. A.L.S. and R.J.L.T., with input from R.O.D., performed and analyzed structural simulations. E.L.A. provided critical reagents. A.L.S. and R.D.K. wrote the manuscript with input from all authors.

## Declaration of interests

The authors declare no competing interests.

## Notes

### Competing Interest Statement

The authors have declared no competing interest.

