## Supplemental figures and methods for "Simple biochemical features underlie transcriptional activation domain diversity and dynamic, fuzzy binding to Mediator"

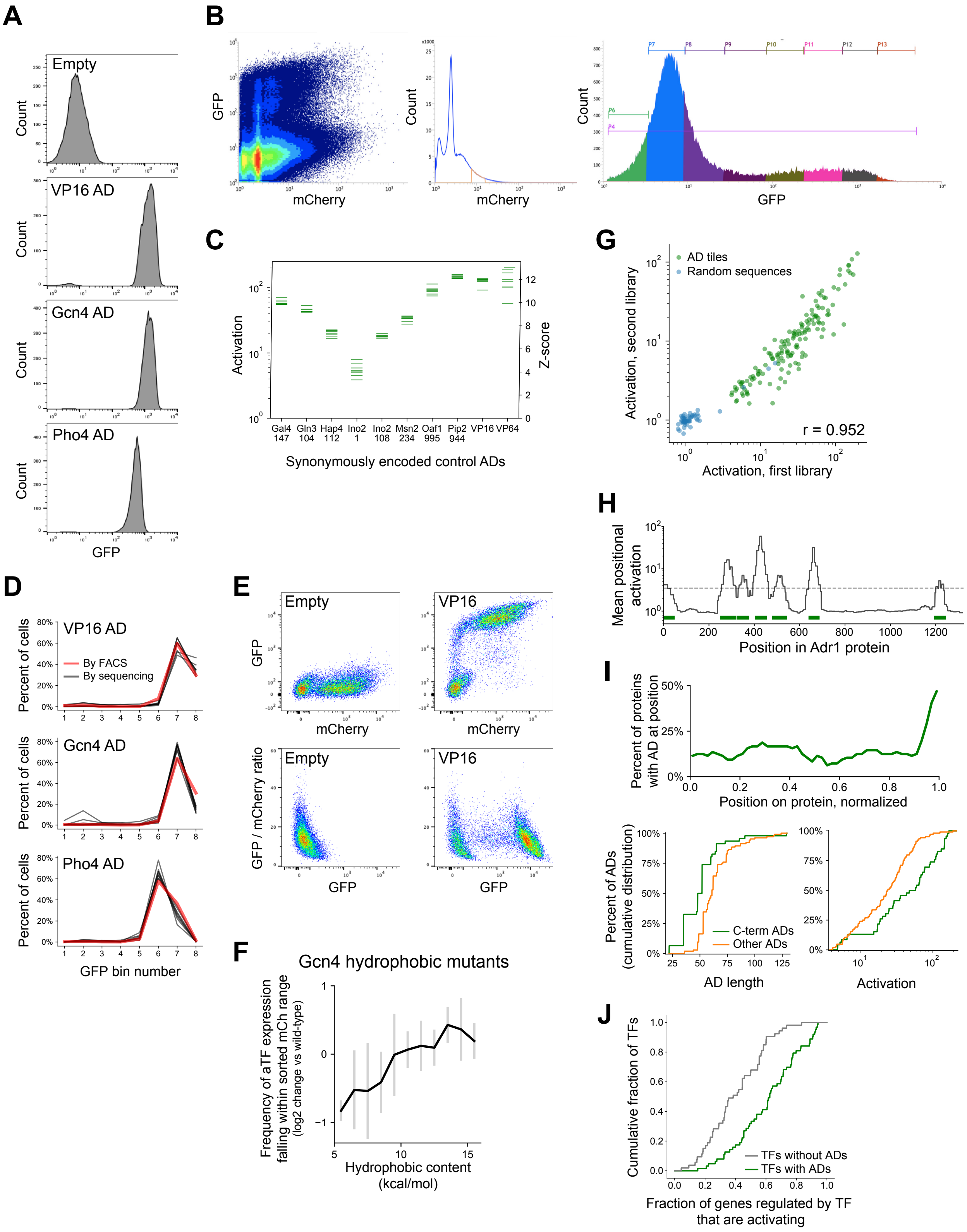


**Figure S1. Methodology, validation, and summary statistics for pooled activation screens. Related to Figure 1.**

A. Distributions of signal from the GFP reporter (in arbitrary units), measured from FACS, in cells expressing artificial TF (aTF) fused to ADs from VP16, Gcn4, or Pho4, or without an AD. Cells were filtered by a narrow range of mCherry, which tracks the aTF expression level.
B. FACS plots of 1,000,000 events observed in the activation assay with the TF tiling and AD mutant sub-libraries (A and B), showing (left) mCherry and GFP signal for each cell, (middle) the mCherry range used for sorting, and (right) the distribution across the eight GFP bins.
C. *In vivo* fold-activation and corresponding Z-scores measured for 10 positive control ADs that were each encoded by 8 different synonymous DNA sequences. Horizontal axis labels are protein and start position of each 53-aa AD.
D. Distributions across the 8 GFP bins of cells expressing aTF fused to ADs from VP16, Gcn4, or Pho4, either measured individually by FACS (red line) or in the pooled screen encoded by 8 different synonymous DNA sequences (black lines).
E. mCherry versus GFP and GFP versus GFP/mCherry ratio for cells expressing aTF with no AD or with the VP16 AD. The GFP/mCherry ratio alone poorly distinguishes GFP-positive cells.
F. aTF expression is affected by tile sequence. To estimate how tile protein sequences affected aTF expression levels, we sequenced all cells used as input to FACS and examined, for each sequence, the frequency at which input cells fell within the filtered mCherry range (i.e. were seen across all 8 GFP bins). Plotted here is the mean (black) and standard deviation (grey bars) of this measure for the hydrophobic mutants of the Gcn4 AD (Figure 2F), relative to wild-type, when binned by their hydrophobic content.
G. Scatter plot of *in vivo* activation of tiles from 150 ADs (green) and 50 random sequence controls (blue) shared between two separately cloned and assayed libraries (sub-libraries A and D). Pearson r = 0.952.
H. ADs were annotated using the (log-scale) mean fold-activation at each protein position, shown here for Adr1. Width of ADs were determined by the full width half maximum of each peak that crossed the threshold (dashed line). Annotated ADs are marked in green bars. Related to Figure 1D-E.
I. (*Above*) C-terminal ADs are in nearly half of all AD-containing TFs. The horizontal axis shows protein position normalized so that the N- and C-termini are at 0 and 1 respectively. (*Below*) Cumulative distribution plots of the length and maximal activation (among overlapping tiles) for ADs at the C-terminus of a protein (green) versus other ADs, showing that C-terminal ADs are shorter and stronger on average.
J. TFs that contained ADs upregulated a higher proportion of downstream genes than TFs without ADs, P = 1.5E-7 by KS test. Genes up- or down-regulated by all TFs were annotated by (Hackett et al., 2020).

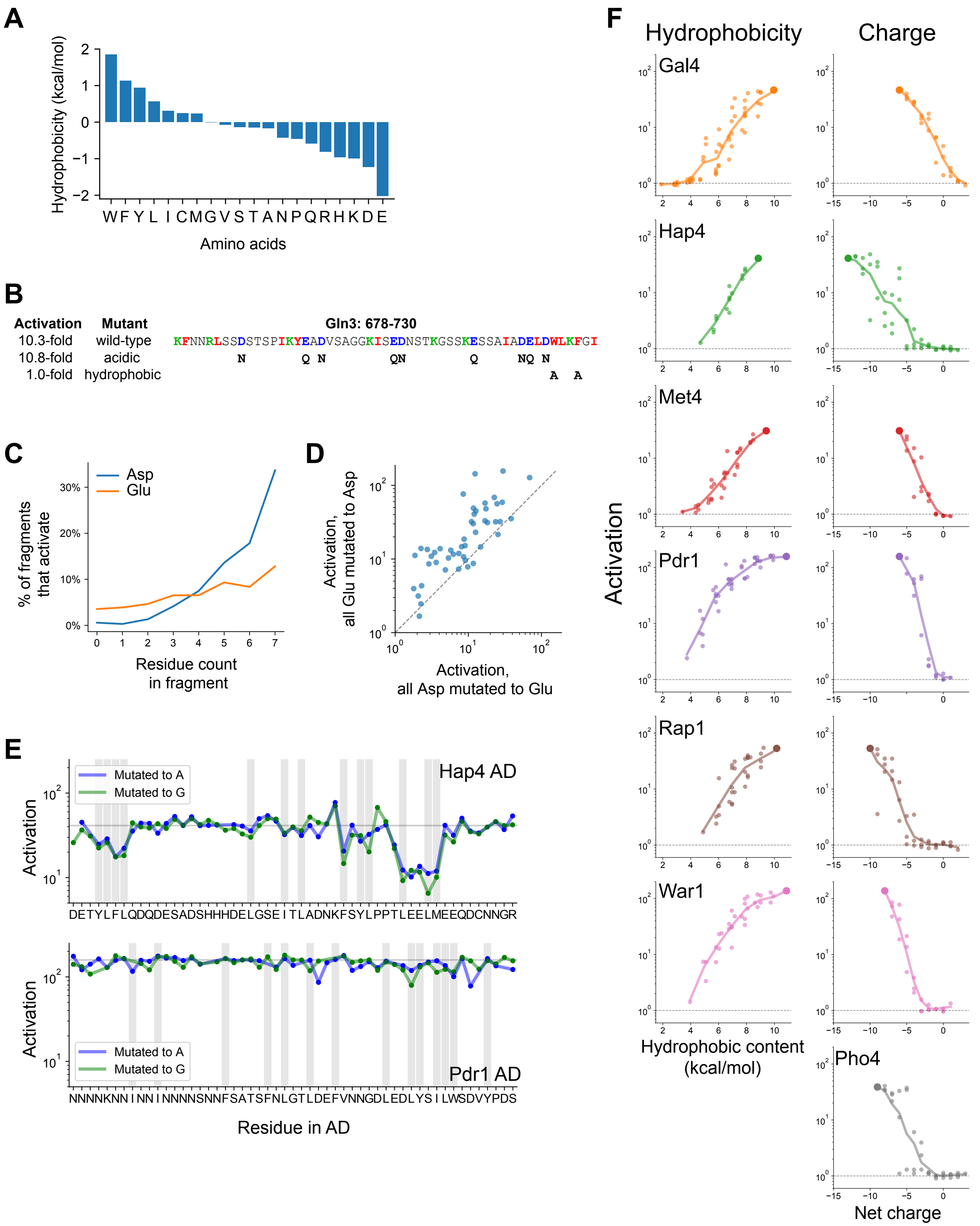


**Figure S2. Mutagenesis of hydrophobic and acidic residues in known and newly-discovered ADs. Related to Figure 2.**

A. The Wimley-White interfacial scale used for calculating hydrophobic content of sequences. Related to Figure 2A-B.
B. The C-terminal AD of Gln3 activated just as strongly at +7 net charge, without acidic residues. However, mutating a pair of aromatic residues prevented activation, identifying the protein C-terminus as a key region. Hydrophobic, acidic, and basic residues are shown in red, blue, and green respectively; mutated residues are shown below the wild-type sequence. Related to Figure 2C.
C. Within fragments that contain exactly 0-7 Asp or Glu residues, the percent that activate ($P < 0.0001$) is plotted. Asp is more strongly associated with activation than Glu.
D. When all Glu residues were mutated to Asp, each of 150 ADs activated comparably to or stronger than when all Asp residues were mutated to Glu.
E. Impact of mutating every individual residue to Ala (blue) or Gly (green) in the Hap4 and Pdr1 ADs. The Hap4 AD has a few specific important residues while residues in the Pdr1 AD are all redundant. Wild-type activation is marked by the horizontal line, positions with bulky hydrophobic residues (WFYILM) are marked in grey. Ala mutants that were within 2-fold of the Gly mutant at the same position were plotted in Figure 2E.
F. All mutants in which aromatic residues (*left*) or acidic residues (*right*) were successively removed, as plotted in Figure 2F. Because the Pho4 AD contained only two aromatic residues, its mutants are not shown.

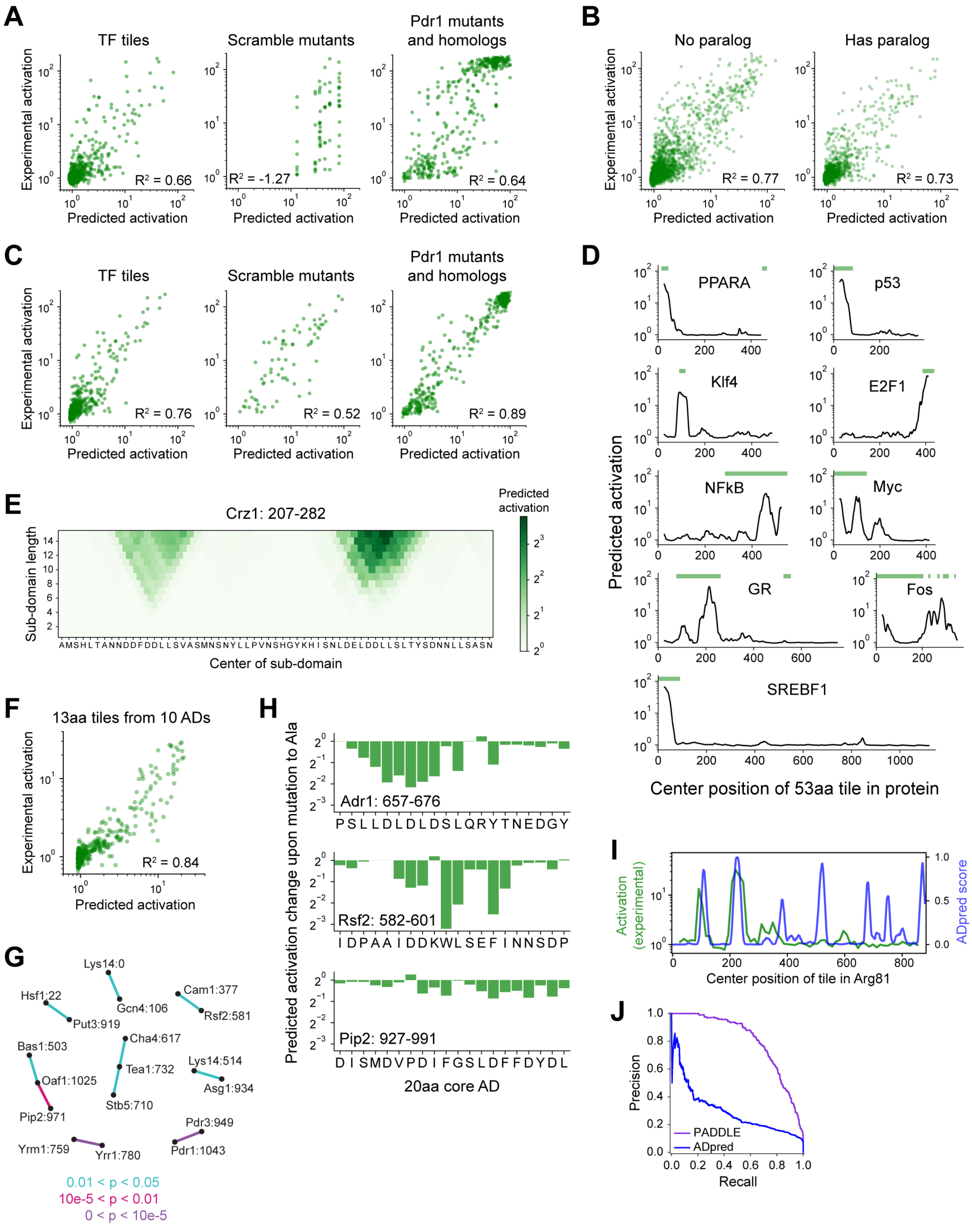

**Figure S3. Validation of neural network models and PADDLE predictions on human TFs and yeast core ADs. Related to Figure 3.**

A. Neural network predictions from amino acid composition alone on test data withheld from training. $R^2$ is the coefficient of determination. Compare to PADDLE predictions in Figure 3C.
B. PADDLE predictions on TF tiles in the training validation set, split by whether the TF has a paralog. $R^2$ is the coefficient of determination. PADDLE is not more accurate on TFs that have a paralog in the training set.
C. Predictions by PADDLE-noSS, which does not use predicted secondary structure or disorder as input, on test data withheld from training. $R^2$ is the coefficient of determination. Compare to Figure 3C.
D. Examples of PADDLE predictions in human TFs compared with previously known ADs (green bars). GR is the glucocorticoid receptor. Related to Figure 3D. AD annotations are from (Barrett et al., 2005; Dowell et al., 1997; Funk et al., 1997; Ghaleb and Yang, 2017; Hi et al., 1999; Kato et al., 1990; Lavery and McEwan, 2005; Raj and Attardi, 2017; Sato et al., 1994; Seipel et al., 1992; Sutherland et al., 1992; Yet et al., 1998).
E. PADDLE-noSS predictions on all sub-domains up to 15 residues in length within positions 207-282 in Crz1. Each sub-domain corresponds to a square and its predicted activation is shown in color. Related to Figure 3E.
F. Scatter plot of predictions and experimentally-measured activation for all 13-aa tiles of 10 ADs. $R^2$ is the coefficient of determination. Related to Figure 3E.
G. The 20-aa core regions from all ADs have no common shared motifs. Pairwise BLAST Expect values (E-values) were calculated between all pairs of core ADs using all-against-all protein BLAST with an Expect threshold of 200,000, an alignment seed ("word") size of 3, BLOSUM62 as the scoring matrix, a gap existence cost of 10, and a gap extension cost of 1. P-values were calculated as 1 - exp(-E). All core AD pairs with P < 0.05 are shown here connected by lines colored according to P-value. Core ADs (nodes) are labeled by their protein and start position.
H. For three 20-aa core ADs, the predicted activation change upon mutating each individual residue to Ala is shown. Related to Figure 3H.
I. Comparison of ADpred (Erijman et al., 2020) predictions on 30-aa tiles in Arg81 (blue), smoothed with a 15-aa moving average, with experimental measurements of activation on 53-aa tiles (green). ADpred correctly predicts two ADs but incorrectly predicts additional ADs. Compare to Figure 3B.
J. Precision-recall curves for predictions by PADDLE (purple) or ADpred (blue). The predictions are evaluated on all wild-type TFs tiles withheld from PADDLE's training dataset. ADpred score for each 53-aa tile is taken as the 80$^{th}$ percentile of ADpred scores for all containing 30-aa tiles. Area under the curve (AUPRC) for PADDLE and ADpred are 0.805 and 0.294 respectively.

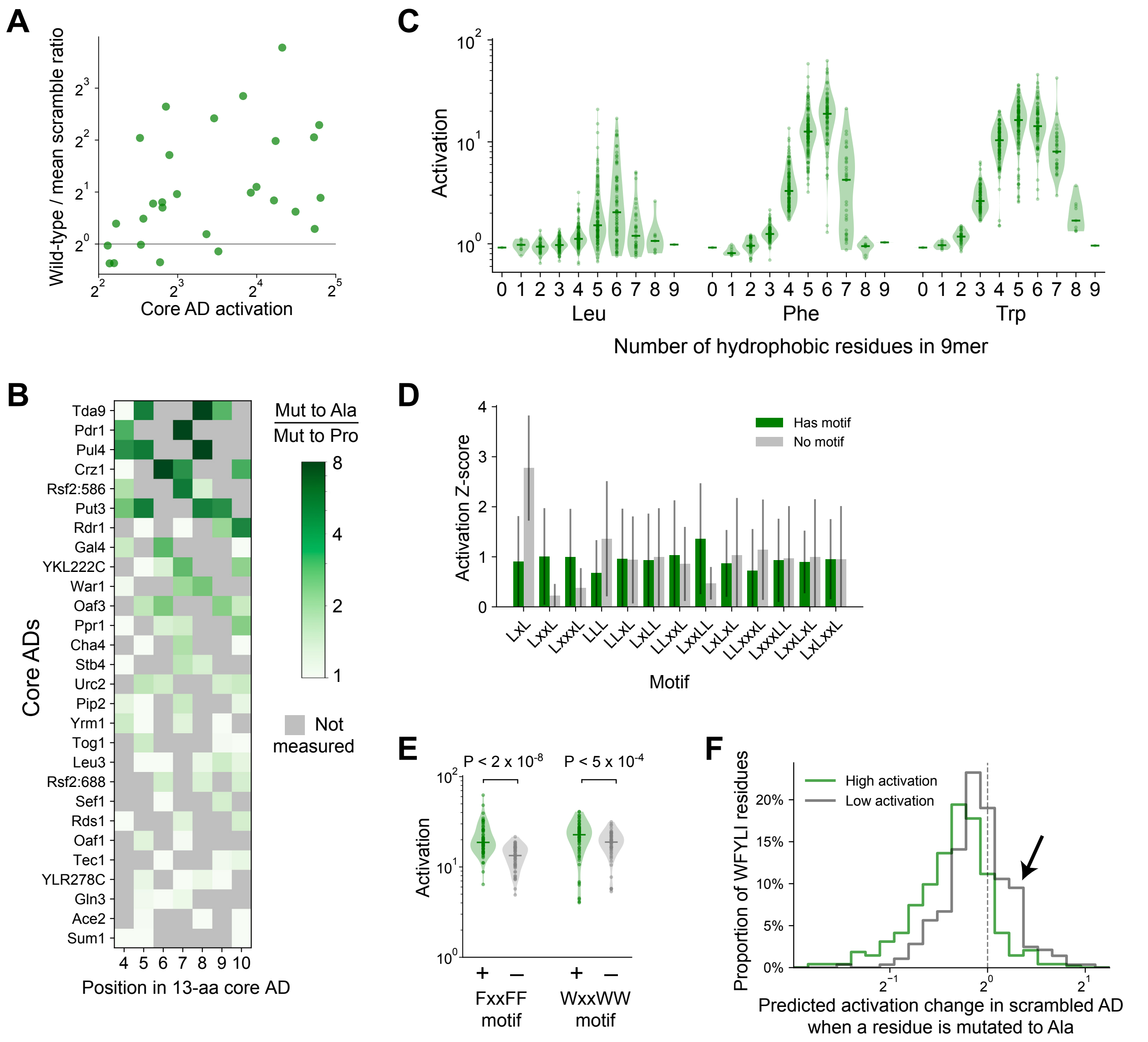


**Figure S4. Additional activation data for and analysis of core AD mutants and systematic 9mers. Related to Figure 4.**

A. The wild-type-to-scramble ratio of core ADs was not correlated with their wild-type activation strength. Related to Figure 4A-B.

B. For each position assayed across all 28 core ADs, the amount that proline disrupts activation—namely the ratio in activation upon mutation to Ala versus to Pro—is plotted in color. Positions with hydrophobic residues were not measured and are marked in grey. The largest ratio observed across different positions is the value plotted for each core AD in Figure 4D.

C. Distribution of activation by all possible 9mers consisting entirely of only two amino acids each—Asp and either Leu, Phe, or Trp—grouped by the number and identity of the hydrophobic residue (i.e. the amino acid composition). Related to Figure 4E-F.

D. Within 9mers containing five Leu residues, the mean and standard deviation activation Z-scores of 9mers with (green) and without (grey) each short motif is plotted. For each motif, P-values were calculated with Kolmogorov-Smirnov test. The LxxLL motif was significant at P = 3.2E-9; the next most significant was LxxL at P = 0.0015. Related to Figure 4G.
E. Within 9mers containing five Phe or five Trp residues, FxxFF and WxxWW motifs were also significant at P = 1.9E-8 and P = 4.8E-4 respectively. Related to Figure 4G.
F. For scramble mutants of the 53-aa ADs (Figure 3A) that activated strongly (P < 0.0001, green) or weakly (grey), we used PADDLE to predict the change in activation upon mutating each individual bulky hydrophobic residue (WFYLI) to Ala. A histogram of these changes is plotted. Weakly-activating scramble mutants have more bulky hydrophobic residues that increase activation when mutated (arrow). Related to Figure 4I-J.

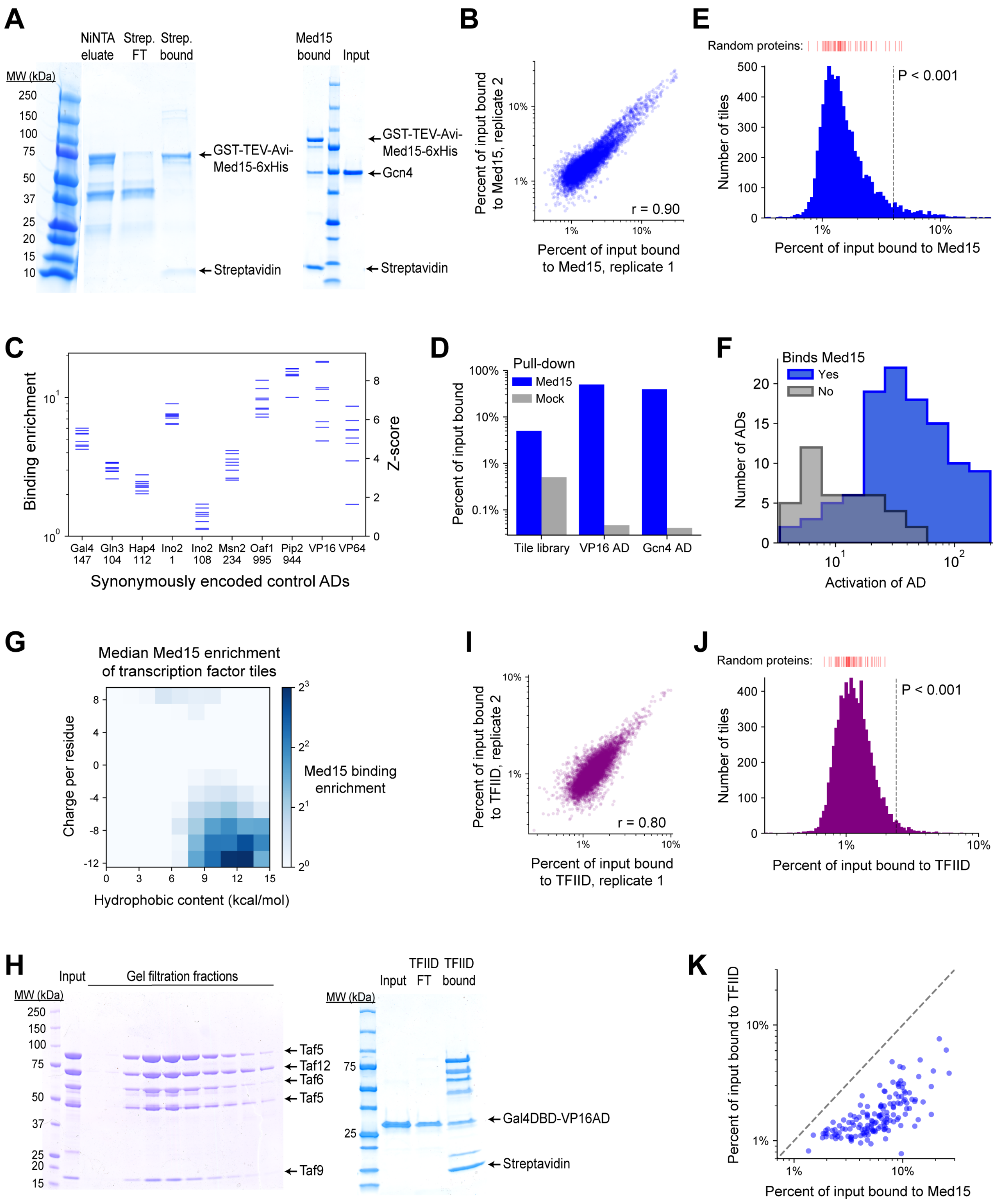


**Figure S5. Methodology, validation, and summary statistics for Med15 and TFIID subcomplex pull-down screens. Related to Figure 5.**

A. (*Left*) A 4-20% polyacrylamide gel showing *in vivo* biotinylated GST-TEV-Avi-Med15K123-6xHis protein (denoted Med15) eluted from NiNTA resin (lane 2) and the flow-through (lane 3) and bound beads (lane 4) after incubation with Dynabeads MyOne Streptavidin T1 magnetic beads. Med15-bound beads prepared in this way were used for pull-down experiments. (*Right*) Gcn4 binds Med15 *in vitro*. Med15-bound beads were incubated with 2ug Gcn4 input (lane 3) for 15 minutes in 20mM Tris pH 8.0**,** 100mM KCl, 1mM MgCl2, 0.01% Tween-20, 1mM DTT and washed twice for 5 minutes each. Bead-bound protein, eluted with SDS and loaded onto this 4-20% polyacrylamide gel, is shown in lane 1.

B. Med15 binding by all TF tiles in two replicate pull-down experiments. Pearson's r = 0.90.
C. Med15 binding and corresponding Z-scores measured for 10 positive control ADs that were each encoded by 8 different synonymous DNA sequences. Horizontal axis labels are protein and start position of each 53-aa AD.
D. Percent of input bound to Med15-bound beads (blue) or beads alone ("mock", grey) for mRNA-tagged and pooled protein fragments of VP16 AD, Gcn4 AD, and the TF tiling library. Binding is tracked by qPCR of cDNA tags with primers specific to VP16 and Gcn4 AD sequences or amplifying all TF tiles.
E. Histogram of Med15 binding for all TF tiles. Binding of 50 random protein sequence controls are marked above in red bars. Vertical dashed line shows the cutoff for P < 0.001. Related to Figure 5B.
F. Histograms of AD activation strength for ADs that bind (blue) or do not bind (grey) Med15. ADs that do not bind Med15 tend to activate more weakly. Related to Figure 5C-D.
G. TF tiles are binned by their charge and hydrophobic content per residue and the median Med15 binding enrichment of each bin is displayed in color. Analogous to Figure 2B, which shows *in vivo* activation.
H. (*Left*) IgG-purified TFIID subcomplex (Taf4/12/5/6/9; "input") and fractions resulting from gel filtration with a Superose 6 Increase column. (*Right*) VP16AD binds TFIID subcomplex *in vitro*. TFIID subcomplex-bound streptavidin beads were incubated with 2ug Gal4DBD-VP16AD input (lane 2) for 15 minutes in 20mM Tris pH 8.0, 100mM KCl, 10mM MgCl2, 0.01% Tween-20, 1mM DTT, and the flow-through was collected (lane 3). After a brief wash, bead-bound protein was eluted with SDS (lane 4).
I. TFIID subcomplex binding by all TF tiles in two replicate pull-down experiments. Pearson's r = 0.80.
J. Histogram of TFIID subcomplex binding for all TF tiles. Binding of 50 random protein sequence controls are marked above in red bars. Vertical dashed line shows the cutoff for P < 0.001. Related to Figure 5B.
K. For each AD, the maximal Med15-binding of all overlapping tiles is plotted against the maximal TFIID subcomplex-binding of all overlapping tiles. (At least 26 aa of overlap is required.) Binding values are percent of input. The dashed line shows equality.

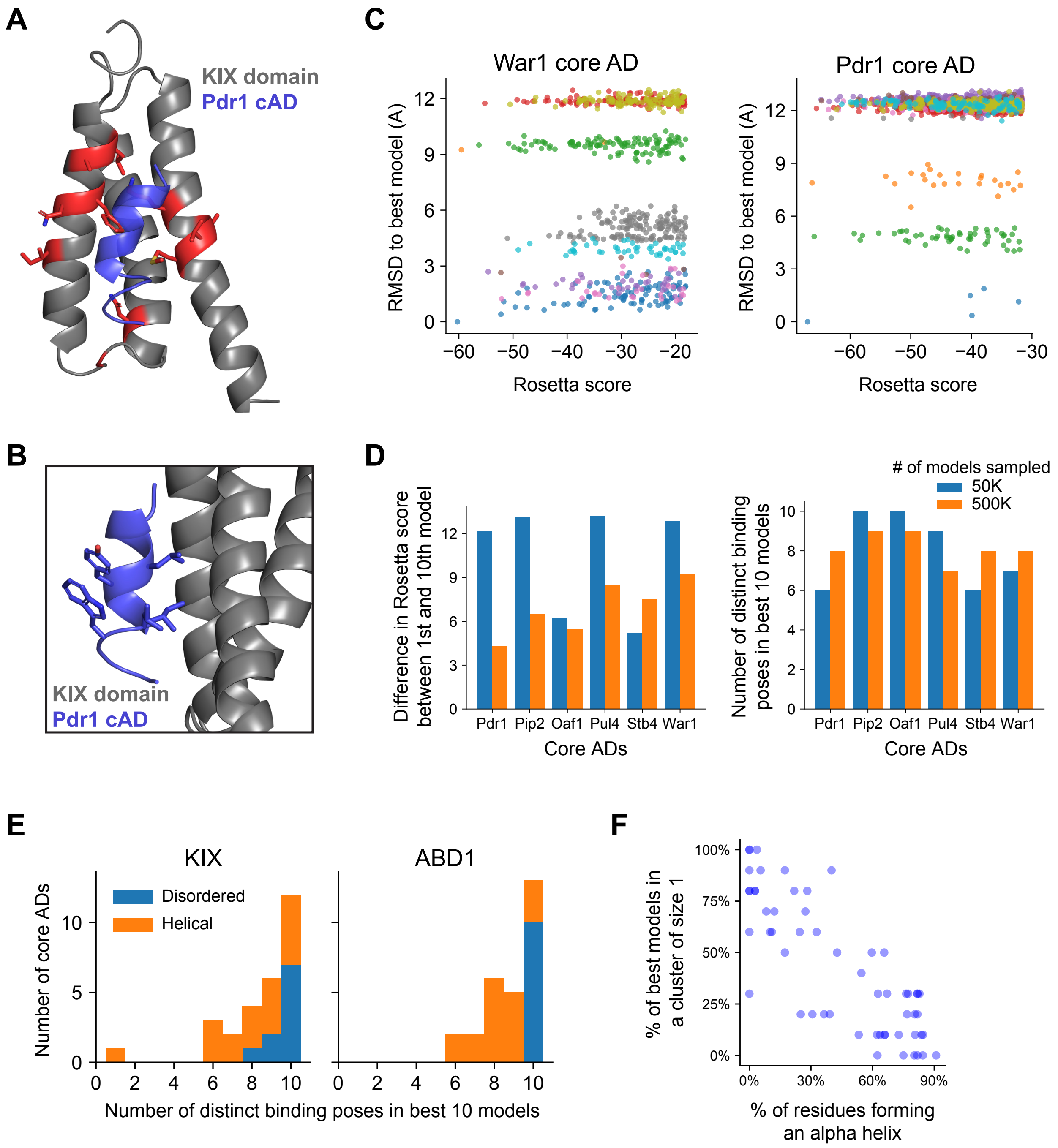


**Figure S6. Validation and additional analysis for structural modeling of core ADs interaction with Med15 ABDs. Related to Figure 6.**

A. The best-scoring model of the Pdr1 core AD (blue) bound to the KIX domain (grey). KIX residues important for binding the Pdr1 AD are shown in red (Thakur et al., 2008). The core AD contacts the hydrophobic surface formed by KIX residues F52, V76, and M77 and forms a salt bridge with K73.

B. The interaction from Panel A shown from the side. Core AD residues L5, I8, and L9 contact the KIX hydrophobic surface but Y6 and W10 are left exposed to solvent. Related to Figure 6C.
C. From the best 5000 out of 500,000 models sampled for the War1 and Pdr1 core ADs interacting with the KIX domain, models from 10 clusters are shown, with each cluster in a unique color. Smaller (more negative) Rosetta scores are better, and the best model is the lower left point. The vertical spread shows the root mean squared distance (RMSD) between the best model and models from other clusters, in Angstroms. Related to Figure 6D.
D. For each of six core ADs binding the KIX domain, a comparison of the 10 best models found when 50,000 (blue) or 500,000 (orange) models were sampled. (*Left*) Sampling ten-fold more models usually reduced the difference in Rosetta score between the 1$^{st}$ and 10$^{th}$ best models. (*Right*) Six or more distinct binding poses (clusters) were observed in the 10 best models even when 500,000 models were sampled. Related to Figure 6E.
E. Core ADs that bound KIX and ABD1 in disordered conformations (fewer than 3 helical residues on average, blue) showed especially high diversity of binding poses. Same as in Figure 6E, but colored by core AD secondary structure and shown as a stacked histogram.
F. In core ADs that had few helical residues when binding KIX or ABD1, most of the 10 best models were the only one their own cluster; i.e. those binding poses were sampled only once.

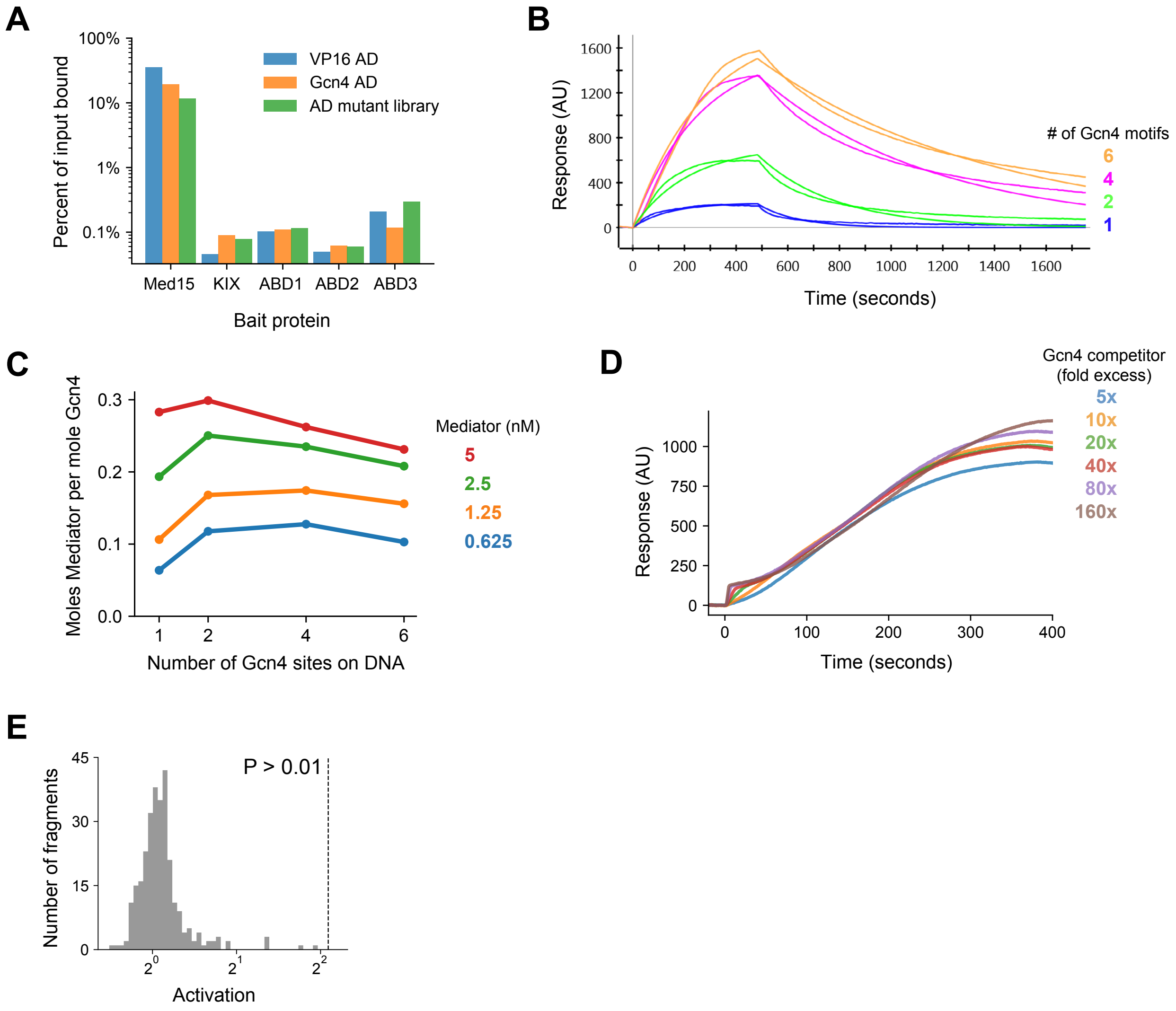


**Figure S7. Additional data and analysis related to Figure 7.**

A. Percent of input bound to Med15 or individual ABDs for mRNA-tagged and pooled protein fragments of VP16 AD, Gcn4 AD, and the AD mutant library (sub-library B). Binding is tracked by qPCR of cDNA tags with primers specific to VP16 and Gcn4 AD sequences or amplifying all AD mutants. Binding to each ABD alone was two orders of magnitude lower than to the four ABDs in tandem.
B. Mediator at 1.25 nM binding to and dissociating from Gcn4-DNA complexes in surface plasmon resonance (SPR) experiments, fit to a Langmuir kinetic model. DNA templates had 1, 2, 4, or 6 Gcn4 motifs. Related to Figure 7C-D.
C. Mediator is recruited more efficiently by Gcn4 on DNA templates with 2 or 4 Gcn4 motifs. The molar ratio of Mediator and Gcn4 bound at equilibrium was determined from their SPR response (in arbitrary units) normalized by their relative mass. Related to Figure 7C-D.
D. Gcn4 competition assay. Binding of 1nM Mediator to Gcn4-DNA complexes on the surface was minimally affected by up to 160-fold molar excess of Gcn4 (160 nM). Gcn4 was pre-bound to DNA, which had two Gcn4 motifs, before the Mediator binding shown

here. The small bump at the onset of binding corresponds to rapid, increased Gcn4 binding at high concentrations. Related to Figure 7E.

E. Histogram of *in vivo* activation for 291 control tiles in yeast nuclear proteins that had extremely high acidic and hydrophobic content but were predicted by PADDLE to not activate. Specifically, these tiles cover all regions in nuclear proteins that have net charge at most -8 and hydrophobic content at least 10 kcal/mol (compare to Fig 2A-B) but predicted activation Z-score less than 0.5. No tiles activated significantly. Related to Figure 7G.

**Table S1.** Data from activation assays. Related to Figures 1-5 and 7.

**Table S2.** ADs identified in the activation screen and their overlap with previously-known ADs. Related to Figure 1. Protein positions are 0-indexed and inclusive of start but exclusive of stop positions.

**Table S3.** ADs predicted in human and virus proteins and core ADs predicted in yeast. Related to Figures 3 and 4. Protein positions are 0-indexed and inclusive of start but exclusive of stop positions.

**Table S4.** Data from pull-down assays and list of Med15-binding domains. Related to Figure 5.

**Table S5.** All FlexPepDock runs. Related to Figure 6.

### Methods

#### RESOURCE AVAILABILITY

#### EXPERIMENTAL MODEL AND SUBJECT DETAILS

The *S. cerevisiae* strain BY4711 (ATCC 200873) with genotype *MATalpha trp1delta63* was used for activation experiments. Standard cell growth conditions were 30C incubation in YPD medium (1% w/v Yeast Extract, 2% w/v Peptone, 2% w/v Dextrose) or synthetic complete (SC) medium (Dunham, 2015). Growth in SC medium lacking tryptophan (SC-Trp) was used to select for transformants with the pRS414 plasmid backbone. Selection for stable transformants harboring the *natMX6* marker gene was performed by supplementing media with 100 µg/mL nourseothricin (NAT). Selection for stable transformants harboring the Kan*MX* marker gene was performed by supplementing media with 200 µg/mL geneticin (G418). In activation assays, overnight cultures were diluted to OD600 of 0.4 in SC-Trp-NAT-G418 with 10 nM beta-estradiol and grown for 4 hours at 30C before FACS.

HEK293T cells (ATCC CRL-3216) were cultured in DMEM with high glucose and GlutaMAX (Thermo 10566016) supplemented with 10% fetal bovine serum (Millipore TMS-013-B) at 37C in 5% $CO_2$.

#### METHOD DETAILS

##### Activation screen plasmid library construction

The artificial TF (aTF) expression cassette, which contains a KanMX gene and an ACT1 promoter driving expression of an aTF (comprising an mCherry tag, mouse Zif269 DBD, and estrogen binding domain) with a multiple cloning site for insertion of protein fragments of interest, was derived from pMVS142_pACT1_mCherry_Zif268_EBD_MCS_KAN, which was a gift from Barak Cohen (Addgene plasmid # 99049) (Staller et al., 2018). This cassette was cloned into the pRS414 backbone, which contains a *TRP1* marker (Sikorski and Hieter, 1989). pRS414 plasmid was digested with EcoRI-HF and SacI, the aTF cassette was PCR-amplified using Phusion polymerase (NEB), and fragments were assembled through Gibson assembly and transformed into XL10-Gold Ultracompetent Cells (Agilent). The final construct, called pRS414-TFchassis, was confirmed by Sanger sequencing.

Oligo pools encoding all the protein fragments to be tested in the screen were synthesized at 12K scale (Twist Bioscience). Each oligo was 200 nt in length and contained a 20-nt 5’ constant region and a 21-nt 3’ constant region which were used for PCR amplification. Different sub-libraries had different 3’ constant regions, which allowed specific amplification of the sub-library from the oligo pool. To enable cloning of the oligo pool into pRS414-TFchassis, 20-bp Gibson homology arms matching pRS414-TFchassis were added using PCR. To amplify each sub-library, 2.5 ng of oligo template was PCR-amplified for 9 cycles using Phusion polymerase, followed by column cleanup. pRS414-TFchassis was digested with NheI-HF and AscI, the amplified oligo pool was cloned in by Gibson assembly, and the product was transformed in two replicates, each into 100 µL of Agilent XL10-Gold cells with 5 µL of Gibson mix. Transformed cells were plated on LB Carbenicillin (Carb) plates to estimate efficiency and also grown in 50 mL LB Carb overnight at 37C for next-day plasmid extraction using Qiagen Midiprep kit.

To generate positive control plasmids for the activation screen, IDT gBlocks containing genes expressing 53-aa-long regions from VP16, Gcn4, and Pho4 ADs were also cloned in the same manner as above. The uncloned pRS414-TFchassis, which lacks an introduced insert, was used as a negative control.

**GFP reporter cell line construction**

pMVS102_P3-GFP-NATMX6 was a gift from Barak Cohen (Addgene plasmid # 99048) (Staller et al., 2018). To construct a yeast GFP reporter cell line, a cassette from pMVS102_P3-GFP-NATMX6 comprising a *natMX6* marker and a P3 promoter (a modified *GAL1* promoter with six copies of the Zif268 binding site) that drives expression of a fast maturing EGFP variant (McIsaac et al., 2013), was PCR-amplified using Phusion polymerase. PCR primers were designed to add 40-bp overhangs with homology to the dubious ORF *YBR032W*. The PCR product was transformed into yeast using a high-efficiency method (Gietz et al., 1992). After overnight outgrowth at 30C, transformed cells were plated onto YPD-NAT, colonies were picked, and integration into *YBR032W* was confirmed by PCR amplification of the integrant and Sanger sequencing.

**Multiple plasmid transformant outgrowth assay**

The *in vivo* activation screen requires a yeast library in which each cell contains one pRS414-TFchassis plasmid with a unique insert. The plasmid is maintained at single-copy or low levels due to its CEN/ARS origin of replication, so if a cell receives multiple different vectors upon transformation then repeated cell divisions will eventually separate the distinct vectors (Scanlon et al., 2009). To ensure generation of a cellular library in which a unique pRS414-TFchassis plasmid was present in each cell, an assay was developed to quantify the fraction of cells initially receiving multiple plasmids and the time required for cells to reach single-copy plasmid levels by repeated cell divisions. First, using a high efficiency transformation protocol (Gietz et al., 1992), approximately 50 million yeast cells in a volume of 360 µL were transformed with 2 µg total plasmid DNA, consisting of an equal mixture of aTF plasmids expressing one of VP16, Gcn4, or Pho4 ADs. Transformed cells were sparsely plated on SC-Trp-NAT plates either immediately or after a 24-hr or 48-hr outgrowth in SC-Trp-NAT liquid media. For each plating condition, colonies were picked for DNA extraction, followed by qPCR to measure the abundance of VP16, Gcn4, and Pho4 plasmids. After 0hr, 24hr, and 48hr of growth, 2 of 15, 2 of 15, and 1 of 24 colonies had multiple vectors present, respectively. Because this method cannot detect multiple transformation events of the same plasmid sequence in the same cell (e.g. double VP16 transformation), a correction was applied, yielding multiple vector rates of 20%, 20%, and 6% for each outgrowth condition, respectively.

**Pooled activation assay in yeast**

The cloned pRS414-TFchassis plasmid library was transformed into the GFP reporter yeast strain using the same method as in the multiple transformant outgrowth assay above but scaled up 2.5-fold. To generate clonal control strains, cells were also separately transformed with a positive control plasmid expressing a VP16, Gcn4, or Pho4 AD or with a negative control plasmid, the pRS414-TFchassis plasmid without a protein fragment insert. Transformed cells were grown in 200 mL SC-Trp-NAT-G418 for 6 days (first library) or 5 days (second library); cells were diluted down to OD = 0.05 every day upon reaching saturation. After this period, cells were diluted to OD600 = 0.4 in SC-Trp-NAT-G418 containing 10 nM beta-estradiol and grown at 30C for 4 hours. Cells were then pelleted and washed once in FACS buffer (PBS + 2mM EDTA + 0.1% BSA, sterile filtered), and finally suspended in FACS buffer at roughly 40 million cells/mL.

Cells were analyzed and sorted on a BD Influx Cell Sorter at the Stanford Shared FACS Facility. Cells from different sub-libraries were pooled just prior to sorting. The largest 10-15% of cells were excluded from sorting, and then an additional filter was applied to limit mCherry signal to within a two-fold range. For the remaining cells, the full range of GFP signal was divided into 8 evenly spaced bins in log scale; cells were first sorted into the 4 low bins and then sorted into the 4 high bins. In total, 2.85 million cells and 2.65 million cells were sorted for the first and second libraries, respectively.

To define the background GFP signal, cells containing the negative control plasmid were also induced with 10 nM beta-estradiol and their distribution across the 8 GFP bins defined above was measured. We found that these cells had slightly increased levels of GFP signal relative to untransformed cells, potentially because binding of the aTF alone slightly opens up chromatin around the GFP promoter. Cells expressing just one plasmid containing the VP16, Gcn4, or Pho4 AD were also individually analyzed (by similarly measuring their distribution across same 8 GFP bins) and compared to results obtained for identical sequences in the pooled activation screen.

Cells sorted into each of the 8 GFP bins and a sample of total input cells (9 samples total) were grown overnight and then pelleted. Pellets were treated with lyticase and then plasmid DNA was extracted using a Qiagen Miniprep kit. The region of the plasmid encoding the variable protein fragment was PCR-amplified for 11-20 cycles (based on the OD600 of the culture) using Phusion polymerase with PCR primers that added flanking sequences corresponding to the standard Illumina sequencing primers. After a column cleanup, a second PCR was performed with primers that added sample-specific barcodes and the P5 and P7 sequences for Illumina sequencing. This second PCR product was SPRI size-selected using AMPure XP beads (Beckman Coulter) at 1x dilution, and the different samples were pooled for paired-end sequencing on the Illumina NextSeq 550. Roughly 2 million reads of 2 x 76 bp length were sequenced per sample.

**Activation assay in human cells**

The pFN26A BIND plasmid (Promega), which contains a Renilla luciferase gene used for normalization, was used to express domains of interest fused to a GAL4 DBD. Genes expressing 50 randomly chosen putative human ADs predicted by PADDLE were synthesized in a single oPool (IDT) with Gibson homology arms, amplified using Phusion polymerase, and cloned into pFN26A BIND (digested with AsiSI and PmeI) using Gibson assembly. The product was transformed into Agilent XL10 Gold cells, and then plated onto LB Carb plates. Forty-three colonies were picked, which yielded clones of 25 correct unique ADs. In addition, plasmids expressing a VP16 AD positive control or three random protein sequences for negative controls were individually cloned in the same manner.

HEK293T cells were plated at 24,000 cells per well of a 96-well plate. The next day, cells were transfected using Lipofectamine 3000 with 50 ng of cloned pFN26A BIND plasmid and 50 ng of pGL4.35 (Promega), which contains a Firefly luciferase reporter gene under the control of a minimal promoter driven by 9 upstream GAL4 DNA binding sites. Each plasmid was transfected into triplicate wells. Seventy-two hours after transfection, luminescence from renilla and firefly luciferases were measured using the Dual-Glo Luciferase assay system (Promega) on a BioTek Synergy 2 plate reader. Measured activity of each predicted AD or control sequence was computed by dividing the Firefly luciferase signal by the Renilla signal and then averaging the triplicate values. Fold-activation was computed by dividing the activity of each predicted AD by the mean activity of the three negative controls, Z-scores were computed based on log-scale fold-activation by standardizing the negative controls to mean of 0 and variance of 1, and P-values were computed with one tailed Z-tests.

**Cloning and purification of proteins**

**Med15.** *S. cerevisiae* Med15 contains four ABDs: KIX (residues 6-91), ABD1 (residues 158-238), ABD2 (residues 277-368), and ABD3 (residues 484-651) (Herbig et al., 2010). We purified the N-terminal portion of Med15 containing the four ABDs, termed Med15K123 and consisting of wild-type residues 1-238, 273-372, and 484-651, as well as each of the four individual Med15 ABDs. Each was cloned and purified with a C-terminal 6xHis tag and a N-terminal Glutathione S-transferase tag, TEV cleavage site, and AviTag (e.g. GST-TEV-Avi-Med15K123-6xHis). DNA expressing Med15K123 and the GST-TEV-Avi tag fusion was codon optimized, synthesized as gBlocks from IDT, and cloned into a pET-28a vector digested with NotI-HF and NcoI-HF (NEB) using Gibson assembly. Individual ABDs were also PCR-amplified from the Med15K123 gBlock and then cloned similarly. Expression plasmids, together with pBirAcm (pACYC184, Avidity), which expresses BirA enzyme for *in vivo* biotinylation, were transformed into BL21 Star (DE3) cells (Thermo) and cultured in LB media with kanamycin and chloramphenicol, supplemented with 50 µM biotin. Expression was induced by addition of 0.5 mM IPTG, and cells were grown overnight at 16C. Cell pellets were suspended in wash buffer (50 mM Tris pH 8.0, 100mM NaCl, 1mM EDTA), pelleted, and then resuspended in lysis buffer (50 mM HEPES pH 7.0 at 4C, 500 mM NaCl, 10% glycerol, 40 mM imidazole, 1 mM DTT, 0.2 mM PMSF, 2 mM benzamidine, 1 mg/mL pepstatin A, 2.5 mg/mL leupeptin). Cells were lysed by passaging twice through the Avestin Emulsiflex C3 (ATA Scientific) at 15,000 psi. The extract was clarified by centrifugation at 40,000 rpm in a T647.5 rotor for 30 min. Clarified extract was bound to Ni-NTA Superflow resin (Qiagen) for 30 min at 4C with rotation, washed with 20 column volumes of lysis buffer, and eluted in 2 column volumes of elution buffer (composed of lysis buffer with 500 mM imidazole). Further enrichment of the full-length purified protein was achieved by binding to streptavidin beads and washing three times before use in pull-down experiments.

**TFIID subcomplex.** The open reading frames of *TAF4*, *TAF5*, *TAF6*, *TAF9* and *TAF12* were PCR-amplified from *S. cerevisiae* genomic DNA (BY4741) and individually cloned into pFASTBac1 (ThermoFisher). *TAF12* was further modified to include an N-terminal 2xProtA tag followed by an AviTag for biotinylation. These tags were separated by a TEV protease cleavage site. BirA, derived from pBirAcm (Avidity), was also PCR-amplified and cloned into pFASTBac1. All plasmids constructs were sequence verified. For protein expression, recombinant baculoviruses were generated per manufacturer's instruction using the Bac-to-Bac expression system (ThermoFisher). The 5-Taf complex, composed of Taf4, Taf5, Taf6, Taf9 and 2xProtA-TEV-Avi-Taf12, was expressed via viral co-infection of Sf9 cells (Expression Systems) with recombinant baculovirus, using an estimated MOI of 5. Taf12 was biotinylated *in vivo*, also via co-infection with a recombinant baculovirus containing BirA. For expression, Sf9 cells were grown at 28C in fernbach flasks shaking at 110 rpm in ESF921 (Expression Systems) supplemented with 50 mM D-biotin for 52-55 hours. Cell pellets were resuspended in lysis buffer (100 mM Tris-Acetate pH 7.9, 5 mM $MgSO_4$, 1 mM EDTA, 195 mM $(NH_4)_2SO_4$, 10% glycerol, 2 mM DTT, 0.2 mM PMSF, 2 mM benzamidine, 1 mg/mL pepstatin A and 2.5 mg/mL leupeptin). Cells were processed for purification immediately or frozen in liquid nitrogen and stored at -80°C. For purification, the resuspended cells were lysed by sonication, followed by centrifugation at 175,000 x g for 1 hr. The clarified extract was passed over IgG Sepharose, in column format, and washed with 5 column volumes of lysis buffer and 5 column volumes of wash buffer (20 mM HEPES-KOH pH 7.9, 75 mM ammonium sulfate, 1 mM $MgSO_4$, 1 mM DTT). The protein complex was eluted with 1.2 column volumes of TEV cleavage buffer (wash buffer supplemented with 15 µg/mL TEV protease) for 12-16 hours at 4C. The TEV elution was subsequently concentrated using a Vivaspin 20 column (PES membrane, 100-kDa molecular weight cut-off) and applied to a Superose 6 Increase gel filtration column, normalized with wash buffer + 5% glycerol. Peak fractions were pooled, concentrated to 4.26 mg/mL and flash frozen in liquid nitrogen.

**Mediator complex and GCN4**. *S. cerevisiae* Mediator complex used in surface plasmon resonance experiments was purified as in (Robinson et al., 2012), except that DNA removal was achieved by adding polyethylemine (PEI) to the cell extract to a final concentration of 0.2%, followed by ammonium sulfate precipitation. Gcn4 was expressed in a pET30a vector and purified also as described in (Robinson et al., 2012).

**Gcn4AD-EcoRI(E111Q)**. Residues 1-224 of Gcn4, corresponding to the full protein except the DNA-binding domain, was PCR-amplified from *S. cerevisiae* genomic DNA. The gene for a nuclease-deficient EcoRI with mutation E111Q was synthesized as a gBlock from IDT and then PCR amplified (Wright et al., 1989). Both PCR products were cloned into a pET-28a plasmid that was NotI-HF and NcoI-HF digested using Gibson assembly. Gcn4AD-EcoRI(E111Q)-6xHis was purified in the same manner as the Med15 proteins, except using a different lysis buffer (50 mM Tris pH 7.5, 500 mM NaCl, 10 mM beta-mercaptoethanol (bME), 20 mM imidazole, 5% glycerol, protease inhibitors) and different elution buffer (20 mM Tris pH 7.5, 300 mM NaCl, 0.1% triton X-100, 10 mM bME, 300 mM imidazole, 5% glycerol).

**Expression of mRNA display libraries**

An mRNA display library was made for each of sub-library A, sub-library B, and 6 pooled controls (3 libraries total), using the following method. Sub-libraries of wild-type and mutant TF fragments were expressed as mRNA-tagged peptides using mRNA display with *in vitro* translation as described in (Takahashi and Roberts, 2009). Specifically, sub-library pools were PCR-amplified using primers that added an upstream T7 promoter sequence and a downstream linker and HA-tag but no stop codon. Thus, all encoded peptides were of the form

MSGT[variable protein sequence]TSVGGSGSYPYDVPDYA and fused to mRNA at the C-terminus. In addition to the two sub-libraries, similar DNA sequences encoding positive control ADs from VP16, Gcn4, and Pho4 and three negative control random protein sequences were synthesized as gBlocks (IDT) (containing an upstream T7 promoter sequence and a downstream linker and HA-tag but no stop codon).

All subsequent work was performed in RNase-free conditions. mRNA was *in vitro* transcribed for 4 hours at 37C using the MEGAscript T7 transcription kit (Thermo) and purified with RNeasy mini columns (Qiagen). The PF30P oligo containing a poly-A sequence, spacers, and puromycin was synthesized by IDT and ligated to the 3' end of the mRNA using Moore-sharp splinted ligation by T4 DNA ligase. The ligation product was desalted by three rounds of concentrating in 0.5 mL Amicon Ultracel 30K MWCO columns and diluting in water. The desalted product was run on a denaturing 4% polyacrylamide gel with 7 M urea. The band corresponding to full-length ligated mRNA was excised from the gel and recovered using the Model 422 Electro-Eluter with 3500 MWCO caps (Bio Rad). After concentrating and desalting in the same manner, the ligated and purified mRNA was used in an *in vitro* translation reaction using the Retic Lysate IVT kit (Thermo) for 60 min at 30C. $MgCl_2$ and KCl were added to final concentrations of 60 mM and 500 mM, respectively, and the sample was incubated for 15 min at room temperature to encourage puromycin fusion. The sample was then diluted in binding buffer (20 mM Tris pH 8.0, 100 mM KCl, 1 mM $MgCl_2$, 0.01% Tween-20, 0.1 mg/mL BSA, 0.1 mM DTT) and bound to Anti-HA magnetic beads (Pierce) for 30 min at room temperature with mixing. After washing three times for 5 min with binding buffer without BSA, peptides were eluted by incubating in 1x SuperScript II buffer (50 mM Tris-HCl pH 8.3, 75 mM KCl, 3 mM $MgCl_2$; Thermo) with 0.01% Tween-20 and 0.5 mg/mL synthetic HA peptide (Thermo) for 1 hour at room temperature with mixing. The mRNA tag on the fused peptides was reverse transcribed using SuperScript II (Thermo) and a primer that contained (1) a sequence that hybridizes just downstream of variable protein sequence coding region, (2) a 12-nt random sequence that serves as a unique molecular identifier (UMI), and (3) the Illumina read 2 sequencing primer sequence. Enzyme was heat inactivated by adding EDTA to 10 mM and heating to 65C for 10 min. To remove aggregates, the library was fractionated in 20 mM Tris pH 8.0, 100 mM KCl, 1 mM $MgCl_2$, 1 mM DTT over a 6-mL 10-30% sucrose gradient by spinning at 60,000 rpm in a SW60 rotor for 10 hours. Fractions of 100-200 µL were collected and the abundance of peptide-mRNA-cDNA fusion in each fraction was assayed using qPCR. The 3-4 peak fractions encompassing 80-90% of total input was pooled and used in pull-down experiments or flash frozen in aliquots and stored at -80C.

**Pull-down assays**

Biotinylated bait proteins (12 µg of Med15 or 8 µg of TFIID subcomplex) were bound to 50 µL Dynabeads MyOne Streptavidin T1 (Thermo) for 15 min at room temperature in binding buffer (20 mM Tris pH 8.0, 100 mM KCl, 10 mM $MgCl_2$, 0.01% Tween-20, 1 mM DTT) and then washed three times for 5 min in binding buffer. In the third wash, beads were blocked by also including 100 ng/µL salmon sperm DNA (Thermo) and 10 ng/µL BSA (NEB) in the buffer. mRNA display library input was prepared by mixing sub-library A, sub-library B, and the three positive and three negative controls. This input was then incubated with the bait-coated beads in binding buffer with salmon sperm DNA and BSA at 100 µL total volume for 8 min with mixing.

Afterward, the beads were separated on a magnet, briefly washed once with cold binding buffer, and then immediately separated on a magnet again. cDNA from bound fragments was eluted by suspending the beads in 10 mM Tris pH 8.0 with proteinase K (NEB) and incubated for 30 min at room temperature. Proteinase K was heat inactivated at 95C for 10 min, beads were separated on a magnet, and the eluate was collected.

Total abundance of the library cDNA in the input, bound, and eluted samples was measured by qPCR. To measure the total abundance of the whole library (without regard to the identity of the library elements), qPCR primers were designed to hybridize to the constant sequences flanking the variable region of the library. Additionally, this design enabled the use of different primers to separately track sub-library A and sub-library B, as well as each of the six control sequences individually. The controls were used to provide an estimate of binding efficiency. Binding percentage was calculated by dividing the abundance of each library in the bound sample by the abundance in the input sample.

cDNA of the input and bound samples were prepared for sequencing in the same manner as the purified plasmids in the activation assay.

**Surface plasmon resonance (SPR) experiments**

Biotinylated 104-bp double-stranded DNA templates were assembled by annealing a 104-nt oligo to a 20-nt oligo with a 5' biotin modification (IDT) and elongating using Klenow Fragment (3'-->5' exo-) (NEB). The oligos were designed such that the TF DNA binding sites were placed away from the biotinylated end. SPR experiments were performed on the ProteOn XPR36 Surface Plasmon Resonance System using NeutrAvidin-coated NLC Sensor Chips (Bio Rad). Mass of biomolecules bound to the chip surface was measured in arbitrary units (AU) approximately every 1 second. All plotted data were normalized to the first ligand channel and the first analyte channel which were used as negative controls in experiments as described below.

In SPR experiments with GCN4, 104-bp DNA templates with 0, 1, 2, 4, or 6 Gcn4 motifs were bound to the chip along the ligand channels, except for the first ligand channel, which was left without DNA for normalization. Binding buffer for all conditions was 20 mM Tris-HCl (pH 7.5), 100 mM KCl, 10 mM $MgCl_2$, 2 mM EDTA, 10 mM $AmSO_4$, 0.05% Tween-20, with 1 mg/mL BSA (NEB). Bound DNA was washed twice with 1 M NaCl. Gcn4 at 7.5 nM was then pre-bound to DNA for 480 seconds along the analyte channels, except for the first analyte channel, which was left without Gcn4 for normalization. After Gcn4 binding, Gcn4 at 7.5 nM with Mediator at 0, 1.25, 2.5, 5, or 10 nM were flowed across analyte channels 2-6 respectively for 300 seconds, then dissociation was observed for 20 min by flowing in binding buffer. Association, dissociation, and affinity constants of Mediator binding to Gcn4 on DNA were estimated by fitting to a Langmuir kinetic model using the ProteOn Manager Software (Bio Rad). Reported values are the mean and standard deviation across the four different Mediator concentrations. Note that Mediator dissociation rates in conditions with multiple Gcn4 binding sites may be overestimated because it also includes the dissociation of Gcn4 from DNA.

For SPR experiments with EcoRI, biotinylated 60-bp DNA templates containing 0 or 1 EcoRI motif were annealed and elongated as described above, and then bound along the ligand channels and washed with 1 M NaCl. Gcn4AD-EcoRI(E111Q) was bound along analyte channels at 27 nM monomer in 20 mM Tris-HCl (pH 7.5), 10 mM $MgCl_2$, 300 mM NaCl, 10 mM bME, 0.05% Tween-20, 1 mg/mL BSA (higher salt was used to reduce non-specific binding to DNA).

Minimal dissociation was observed after binding. Then, Mediator at 5 nM with competitor Gcn4 protein (in molar excess as specified in Figure 7E) was flowed across the chip for 180 seconds in 20 mM Tris-HCl (pH 7.5), 10 mM $MgCl_2$, 100 mM NaCl, 10 mM $AmSO_4$, 10 mM bME, 0.05% Tween-20, 1 mg/mL BSA.

#### QUANTIFICATION AND STATISTICAL ANALYSIS

##### Protein library design

All protein positions reported in the text are 1-indexed and inclusive. The first library of protein sequences was composed of sub-library A, which contained wild-type tiles spanning all *S. cerevisiae* transcription factors, and sub-library B, which contains mutants of 8 known *S. cerevisiae* ADs. The second library was composed of sub-library D and sub-library E, which contained multiple collections of wild-type, mutant, and designed sequences. Sequences in sub-libraries A, B, and D were 53 amino acids in length and sequences in sub-library E were 30 amino acids in length. Protein and DNA sequences, as well as activation and pull-down data where available, are listed in Tables S1 and S4. Libraries contained additional fragments which were included in experiments but are not reported here.

**Sub-library A.** 7637 fragments total. This peptide library was generated from all 162 yeast (Saccharomyces cerevisiae S288C) transcription factors—i.e. genes annotated with the Gene Ontology term GO:003700 (DNA-binding transcription factor activity) (Ashburner et al., 2000; The Gene Ontology Consortium, 2019)—and also MET4 and HAP4. Each transcription factor protein sequence was fragmented into 53-amino acid tiles with at least 40 amino acid overlap (overlap was adjusted based on the protein length to make the tiling as evenly distributed as possible across any given protein), yielding 7460 unique peptides. A set of 50 random synthetic (i.e., not derived from a known protein) sequences with the same amino acid frequencies as the overall set of yeast transcription factors, as well as a set of 50 peptides derived from non-nuclear proteins (proteins not annotated with the GO term GO:0005634 (nucleus)) were used as negative controls. Additionally, 10 positive control sequences from known ADs, some of which overlapped TF tiles, were included. These control protein sequences were each synonymously encoded by 8 different DNA sequences in this sub-library (see below); the mean activation across the 8 encodings was used in further analysis. Data involving this sub-library is shown in Figures 1C-E, 2A-B, 3B-C, 5B-D, S1C-D, S1G-J, S3A-C, S3I-J, and S5B-K.

**Sub-library B.** 2900 fragments total. Eight 53-residue regions from previously reported ADs were selected for more detailed analysis with mutagenesis. These regions were from Gcn4 (82-134), Gal4 (829-end), Met4 (72-124), War1 (892-end), Hap4 (424-476), Pho4 (58-110), Pdr1 (1016-end), and Rap1 (630-682). The wild-type sequence of the eight regions were synonymously encoded by 8 different DNA sequences. For each region, similar classes of mutants were designed, as follows.

- Scanning mutants: Each position was individually mutated either to alanine or glycine. Used in Figures 2E and S2E.
- Scramble mutants: Eight scrambles of each region were randomly generated. Used in Figures 3A, 3C, S3A, and S3C.

- Aromatic mutants: Up to 10 different subsets of *N* aromatic residues (Trp, Phe, or Tyr) were randomly selected and mutated to alanine, where *N* ranged from 1 to the total number of aromatic residues. Used in Figures 2F, 5E, S2F, S3A, and S3C. Also, 1-4 non-hydrophobic positions were randomly selected to mutate to Trp, Phe, or Leu, in 5 ways each.
- Acidic mutants: Up to 3 different subsets of *N* acidic residues (Asp or Glu) were randomly selected and mutated to alanine, where *N* ranged from 1 to the total number of acidic residues. Used in Figures 2F, 5E, S2F, S3A, and S3C. Also, 1-10 non-hydrophobic positions were randomly selected to mutate to Asp, in 3 ways each.
- Homologs: Proteins from other fungal species homologous to S. cerevisiae (yeast) TFs were found by protein alignment to the NCBI nr (non-redundant protein sequence) database using BLAST. A multiple sequence alignment (MSA) was then constructed for each yeast TF and its homologous fungal proteins using Clustal Omega. The homologous fungal proteins were then filtered for those that had at least one match in the MSA to a known yeast ADs (i.e., the region aligned to the known AD was not entirely spanned by a gap). This yielded 285, 251, and 179 homologous sequences for the Gcn4 (120-134), Pho4 (81-110), and Pdr1 (1044-1068) ADs, respectively. Used in Figures 3C, S3A, and S3C.

Two additional sets of 50 random synthetic sequences and 50 peptides derived from non-nuclear proteins were all also included as negative controls, as described for sub-library A.

**Sub-library D.** 2972 unique fragments total. To assess reproducibility, this sub-library included the strongest activating fragment from each of the 150 ADs identified and the 50 random sequence controls from sub-library A, which are shown in Figure S1G.

- AD mutants: For each fragment from the 150 ADs, mutants were included that (1) mutated all Asp to Asn and all Glu to Gln, (2) mutated all Asp to Glu, or (3) mutated all Glu to Asp. Used in Figures 2C, S2B, and S2D.
- Tandem cADs: Analysis of sub-library A data enabled identification of cADs. Forty-seven pairs of identified cADs that could fit within a 53-aa fragment in their native context were chosen. Fragments for this sub-library were designed such that they each contained one of these pairs of cADs, with their original distance between them preserved. Additionally, the first, second, or both cADs were embedded in neutral sequence consisting of AGSTNQV residues, in a manner that maintained the original distance between the two cADs. Used in Figure 7B.
- Hydrophobic inhibition matrix: A 20-aa variable region was placed 5 residues upstream of the 13-aa Pdr1 cAD and embedded within neutral sequence to reach 53 residues total. Random sequences for the variable region were generated, such that relative amino acid frequencies were representative of those from yeast TFs, except for FEKR and WFYLM residues, whose frequencies were systematically inflated to generate a wider range of net charge and hydrophobicity. Eight fragments fitting into each bin were randomly selected to be included in the library. Used in Figure 4J.
- ADs in yeast nuclear proteins: Regions of non-TF nuclear proteins that were predicted by PADDLE to activate were included. Additionally, highly hydrophobic and acidic regions that were predicted by PADDLE not to activate were also included. All designs involving PADDLE predictions are described in more detail below. Used in Figures 7G and S7E.

**Sub-library E.** 3621 unique fragments total. Fragments in this sub-library were all 30 residues in length. Designed sequences of interest, either 13 aa or 9 aa in length, were located at the center of these fragments. The regions flanking these sequences were constant: ASAVGQQG[AN] (upstream) and [NS]SATQSQSNT (downstream), with brackets indicating additional residues for the 9-aa designed sequences.

- AD tiling: Ten identified ADs were selected (from analysis of sub-library A data), and 13-aa tiles spanning each one at 1-aa spacing were included in this sub-library. Used in Figure 3E and S3F.
- Core AD scrambles: The 28 cADs 13 aa in length with the strongest predicted activation (identified from analysis of sub-library A data) were included. Additionally, 33 different scrambled sequences of each cAD were included. Used in Figures 3A-B, 3D, and S4A.
- Proline mutants: In each of the 28 selected cADs, non-WFYILM residues in positions 4-10 were individually mutated to either Ala or Pro. Used in Figures 4C-D, 6B, and S4B.
- Systematic 9mers: All sequences 9 residues in length consisting of only two amino acids each—Asp and either Leu, Phe, or Trp—were included, totaling $3 \times 2^9 - 2 = 1534$ sequences. Used in Figures 4E-H and S4C-E.

Fifty random sequence negative controls were included; these were the first 30 residues of the controls from sub-library B.

**Design of DNA sequences encoding proteins**

In the reverse translation design process, we aimed to optimize our library DNA fragments for compatibility and consistency with our *in vitro* and *in vivo* assays, as well as with standard RNA-seq protocols. We also sought to build in redundancy for error-correcting reads. In particular, we used the Python package dnachisel 1.4.1 (Zulkower and Rosser, 2020) to optimize the following objectives:

1. Use codons matching the relative frequencies in the rabbit species corresponding to the *in vitro* translation kit. Codon frequencies were pulled from the Codon Usage Database hosted by the Kazusa DNA Research Institute (Nakamura et al., 2000).
2. Target an optimal GC content of 45% at both a local (sliding window of 50) level and a global (entire fragment) level.
3. Avoid repeated subsequences of length 10 or more.
4. Avoid homopolymer runs of 8 or more As, 8 or more Ts, 5 or more Cs, or 5 or more Gs.
5. Avoid adjacently repeated k-mers, specifically 3-peats of 3-mers or 5-peats of 2-mers.

To allow paired-end sequencing to uniquely identify each element, we additionally enforced an edit distance of 6 among the first 48 bases and last 48 bases of any two sequences in the same sublibrary. This was performed in a randomized, brute-force, iterative approach, with each iteration consisting of the following steps:

1. Pairwise edit distances were computed for all sequences in the sublibrary.
2. For any sequence with edit distance less than 6 from another sequence, 2 codons in the first 48 bases and 2 codons in the last 48 bases were randomly selected and changed while respecting the encoded amino acid sequence.
3. Repeat until no changes are needed.

Finally, we verified that all sequences sharing the same sub-library primer (e.g. all sequences in sub-library A, or all sequences in sub-library B) had a paired-end edit distance (sum of edit distance of 5'-most 50 bases and edit distance of 3'-most 50 bases) of at least 6.

**Sequencing read alignment**

The use of sequencing primers unique to each sub-library enabled us to submit samples for sequencing in multiplexed format and accurately assign reads to the correct sublibrary computationally. We further leveraged an edit distance margin built into the library to enable mapping of sequencing reads with a small number of errors.

Sequencing read alignment was performed using custom bash script built on top of existing tools and additional custom scripts. It takes as input arguments the UMI length, the sublibrary sequencing primer, the edit distance threshold for that sublibrary, and raw FASTQ files. For pull-down experiments, unique molecular identifiers (UMI) were extracted from reads and appended to the read names using umi_tools 1.0.0 (Smith et al., 2017). cutadapt 1.18 was used to discard reads without matching paired-end sublibrary sequencing primers and trim the primers in reads with matching primers; the default error tolerance was used (Martin, 2011). bwa-mem 0.7.17-r1188 was used to perform a first-pass alignment of reads to the DNA fragment library (Li, 2013). Imperfectly mapped read pairs (i.e., those without paired read SAM flags of 99 and 147) were re-mapped to the library sequence with minimal edit distance. This was necessary because bwa-mem did not always correctly map paired reads as a pair, a problem most evident in the mutant library with many similar sequences. Pairwise Levenshtein edit distance was computed using the Python package editdistance 0.5.3 (https://github.com/roy-ht/editdistance). Paired reads exceeding the edit distance threshold were discarded using reformat.sh from BBTools 38.61 (https://sourceforge.net/projects/bbmap/). Duplicate reads were identified and deduplicated using utmi_tools 1.0.0. Finally, reads mapped to each DNA library fragment sequence were counted.

**Activation assay data processing**

Calculations of GFP signal and fold-activation required the following inputs:

1. The total number of cells sorted into each bin
2. The total number events assayed when sorting bins 1-4 versus bins 5-8
3. The number of sequencing reads of each GFP bin mapped to each fragment
4. The total number of sequencing reads obtained for each bin
5. The average GFP signal observed in each bin during sorting
6. The average GFP signal and number of cells observed for each bin for a negative control population.

First, to make cell counts comparable between sorting bins 1-4 versus bins 5-8, the number of cells sorted into each GFP bin were normalized based on the number of events assayed in each sort. Second, to estimate the number of cells sorted into each of the 8 GFP bins that expressed a given protein fragment *F*, the total number of cells sorted into each bin was multiplied by the fraction of sequencing reads for that bin that mapped to *F*. Any fragment represented by fewer than 10 cells were excluded from further analysis. Third, the average GFP signal of cells expressing *F* was calculated as a weighted average from the average GFP signal observed in each bin during sorting weighted by the distribution of the number of cells expressing *F* across the 8 bins. Because GFP distributions cells expressing single AD sequences were normally distributed in FACS data when GFP signal (arbitrary units) was plotted in log-scale, all averages of GFP distributions were calculated in log-scale (equivalently, a geometric mean of GFP values). The standard deviation of GFP signal of cells expressing *F* was similarly calculated in log-scale and used as a metric for measurement precision. All averages involving activation reported in the paper are calculated in log-scale (equivalently, a geometric mean). Fourth, the fold-activation of *F* was calculated as the mean GFP signal of *F* divided by the mean GFP signal of negative control cells (see below). Z-scores were calculated by standardizing the GFP signal of negative control cells to mean 0 and variance 1 in log scale and P-values were calculated with one tailed Z-tests.

The mean and standard deviation GFP signal of negative control cells was defined in one of two ways. In the experiment with sub-libraries A and B, a sample of estrogen-induced cells containing only the empty pRS414-TFchassis plasmid was assayed under identical conditions and the mean and standard deviation GFP signal was computed from FACS data. In the experiment with sub-libraries D and E, these values were instead determined internally from the 50 random sequence negative controls in the pooled library. Since a small fraction of random sequences activate, outlier sequences were iteratively excluded if they had GFP signal greater than 3 standard deviations away from the mean, resulting in three sequences excluded from each sub-library. Then mean and standard deviation were computed from the sum of the negative control GFP distributions.

In activation assays with sub-libraries D and E, we noticed that many sequencing reads that were mapped with a small number of errors corresponded to fragments with a very large GFP standard deviation. Upon closer examination of sequencing reads of example fragments, the same sequencing errors were consistently found, which likely corresponded to mutations that occurred *in vivo* that affected function of the protein fragment (e.g. inactivation of a strong AD by a frameshift mutation). We therefore used only reads that aligned perfectly for this experiment. Furthermore, fragments with GFP standard deviation greater than 4.5 were excluded from further analysis.

The following three metrics were tracked: the number of total unique fragments, number of fragments observed in at least one of the eight GFP bins, and number of fragments that passed filtering. For the activation assay with sub-libraries A and B, this was (7637, 7604, 7549) for sub-library A and (2900, 2833, 2813) for sub-library B. The same metrics for the activation assay with sub-libraries D and E was (7733, 7716, 7345) for sub-library D and (4261, 4261, 4252) for sub-library E. These numbers include additional fragments in the peptide libraries that are not reported in this paper.

**Activation domain annotations**

To define ADs across all TFs, at each protein position a mean Z-score was calculated from the Z-scores of all fragments overlapping that position. ADs were defined for all intervals of the protein that had a mean positional Z-score (MPZ) greater than 3.5. Starting with intervals that had the highest maximal MPZ score, the start and stop positions of the AD were defined by the full width half maximum; that is, the positions in the protein at which the MPZ fell below half of the maximal MPZ value in that interval. If the AD defined in this manner includes a new region with a MPZ score higher than the original maximal value, this indicated that the MPZ peak was only small relative to nearby peaks, so the AD was then excluded.

**Activation data analysis**

We compared our ADs with the Induction Dynamics gene Expression Atlas (IDEA) (Hackett et al., 2020), which induced expression of every TF individually and measured "v_inter" parameters that describe the resulting fold-change in the expression of all target genes. We said that a TF activated a gene if the v_inter > 1, and for each TF that regulated at least 5 genes, we calculated the proportion of regulated genes that were activated. We then compared these proportions for TFs that had a strong AD (overlapping a fragment with Z-score at least 6) versus TFs without ADs. Significance was calculated using a Kolmogorov–Smirnov test.

Hydrophobicity measurements were calculated using the Wimley-White interfacial scale, which measures whole-residue ΔG for transfer from water to a water-octanol interface (Wimley and White, 1996). Because the goal was to quantify the total content of hydrophobic residues rather than the net hydrophobicity/hydrophilicity, hydrophobic content of a peptide is computed as the sum of the scores of only the hydrophobic residues (i.e. those with negative ΔG). Net charge was computed by adding the number of Arg and Lys residues and subtracting the number of Asp and Glu residues.

Motif finding using DREME was done through the web portal (http://meme-suite.org/tools/dreme) using default settings and shuffled input sequences as control (Bailey, 2011). Searching for 9aaTAD sequences was done by matching to the "most stringent pattern" [MDENQSTYG]{KRHCGP}[ILVFWM]{KRHCGP}{CGP}{KRHCGP}[ILVFWM][ILVFWMAY]{KRHC} listed at the prediction portal https://www.med.muni.cz/9aaTAD/index.php (Piskacek et al., 2007).

**Comparison with ADpred**

ADpred was installed locally from https://github.com/FredHutch/adpred and predictions were computed for all 30-aa tiles on all yeast TFs (Erijman et al., 2020). ADs predicted by ADpred were defined using the authors' criteria, namely sites with five or more contiguous residues with a score >= 0.8. An ADpred-predicted region was considered correct if it overlapped by at least 5 aa with a significantly activating tile ($P < 0.0001$). With this approach, ADpred achieved a precision and recall of 0.366 and 0.692 respectively in predicting experimental ADs. When ADpred-predicted regions were randomly shuffled 100 times to different proteins and positions, the average precision and recall of these shuffled predictions were 0.120 and 0.295 respectively. Alternatively, to compare ADpred predictions directly to our experimental measurements of activation, a representative ADpred score for each 53-aa TF tile in our library was calculated by taking the 80th percentile of ADpred scores of 30-aa tiles contained within the 53-aa tile. These scores were used for calculating tile-wise precision-recall curves for ADpred predictions (Figure S3J).

**mRNA display pull-down data processing**

The fractional pull-down of each fragment *F* was calculated as (the fraction of the total input library that bound to beads, measured from qPCR) x (the fraction of UMI-deduplicated sequencing reads in the bound sample matching *F*) / (the fraction of UMI-deduplicated sequencing reads in the input sample matching *F*). Fragments with fewer than 30 reads in the input sample or fewer than 5 reads in the bound sample were excluded from downstream analysis. Baseline binding was defined using the 50 random protein control sequences in each library, except the sequences that were outliers in the activation assay were excluded. Specifically, random control outliers were iteratively removed if their activation was more than three standard deviations away from the mean (in log scale), resulting in the exclusion of random control #2, 12, 16, 38 from sub-library A and #17, 18, 21, 22, 31, 43 from sub-library B. Binding enrichment of each fragment was then computed as the fractional pull-down of the fragment divided by the mean fractional pull-down (in log scale) of the random controls. Because fractional pull-down and enrichment were normally distributed in log-scale, all averages involving pull-down or enrichment reported in the paper are calculated in log-scale (equivalently, a geometric mean). Z-scores were computed by standardizing the binding enrichment of random controls to mean 0 and variance 1 in log-scale and P-values were calculated with one tailed Z-tests. Triplicate measurements for Med15 pull-downs and duplicate measurements for TFIID subcomplex pull-downs were combined by averaging the binding enrichment values. A small number of highly positively-charged fragments bound in Med15 pull-downs when the mRNA display library was purified on a sucrose gradient but did not bind when the sucrose gradient was omitted. Suspecting that this binding was artifactual, potentially through non-specific binding to mRNA tags, we excluded from analysis 92 fragments that bound with Z-score between -2.5 and 1 without the sucrose gradient and had a Z-score at least 3.5 higher with the sucrose gradient than without.

**Overlap of ADs and Med15- and TFIID subcomplex-binding domains**
Med15-binding domains were annotated in the same manner as ADs using a mean positional Z-score, but with a cutoff of 2.23. An AD was considered to bind Med15 or TFIID subcomplex if it overlapped a fragment that bound significantly ($P < 0.001$) by at least 26 aa. A Med15-binding domain was considered to activate if it overlapped an AD.

**Neural network training**
Neural network models were trained to predict activation Z-scores. Approximately 15% of sequences from sub-library A and B were randomly held-out as a test set, and the remaining library elements were split into 10 parts for training and cross-validation. Because adjacent TF tiles have significant overlap in sequence, all tiles from each protein were placed into the same train or test split. From sub-library B, all mutants and homologs of the Pdr1 AD and all scramble mutants were held-out in the test set and the remaining sequences were distributed evenly across the training splits. Models were developed in TensorFlow 2.2 (Abadi et al., 2016) and trained using CPUs on Stanford's Sherlock computing cluster. All models used mean squared error as the loss function. Sequence encodings and activation Z-scores were standardized to mean 0 variance 1 for more efficient learning. For each architecture, 10 models were trained on each subset of 9 training splits and validated on the 10th split. The best model architecture was chosen by the mean validation score across the 10 splits on sub-library A sequences only. For final evaluation on the test dataset, the mean prediction across all 10 models was used.

**Predicting activation from amino acid composition**

Protein fragment sequences were each encoded as a single 20-element vector giving the proportions of each of the 20 amino acids. The best architecture, as determined by a grid search over hyperparameters, was a neural with three fully-connected layers of width 40 with Swish activation (Ramachandran et al., 2017), batch normalization, L2 weight penalty of 0.01, and dropout rate of 0.4. The model was trained with an initial learning rate of 0.0001 using the Adam optimizer for up to 300 epochs with two scheduling callbacks: reduction of the learning rate by 5-fold if training loss did not improve for 20 epochs, and early stopping if no improvement on the validation loss (averaged in a 10-epoch window) was observed for 75 epochs, upon which the best model from previous epochs was saved.

**Predicting activation from protein sequence (PADDLE)**

One-hot encodings were used to encode each 53-aa protein fragment sequence as a 53-by-20 matrix. Secondary structure predictions were generated using PSIPRED v. 4.01 without PSI-BLAST (Jones, 1999), and a 53-by-2 matrix of the alpha helix and coil predictions were also used as input. Disorder predictions were generated using IUPred2A (Mészáros et al., 2018), and a 53-by-2 matrix of the disorder predictions in “short” and “long” mode were also used as input. In total, each 53-aa protein fragment was encoded as a 53-by-24 matrix.

The best model architecture, as determined by a grid search over hyperparameters, was a convolutional neural network with 9 convolutional layers with kernel size 10 and channel width 30 (i.e. number of filters) followed by max-pooling along the sequence-length dimension and 2 fully-connected layers of width 20, with Swish activation. For regularization, the L2 weight penalty was 0.001, dropout rate was 0.1, and batch normalization was used. The model was trained with an initial learning rate of 0.001 in the same manner as the amino acid composition-based network, except for up to 500 epochs.

After finding this optimal architecture, the test dataset was distributed across the 10 training datasets and 10 new models were trained, together comprising PADDLE. The mean value of these 10 predictors was used as the final prediction. This model was used for all predictions on yeast TFs and nuclear proteins, human TFs, and virus transcriptional regulators. Because secondary structure prediction was by far the slowest aspect of running PADDLE predictions, we trained a version of PADDLE called PADDLE-noSS which only used the one-hot sequence encoding for input, no secondary structure or disorder predictions. PADDLE-noSS was nearly as accurate as PADDLE (see Figure S3C) and was used for predictions of core ADs and mutant sequences in Figures 4E-H, S3E-H, and S4F.

**PADDLE predictions**

Human TFs were defined by Gene Ontology term GO:003700 (DNA-binding transcription factor activity). Virus transcriptional regulators (vTRs) were taken from Table S1 from (Liu et al., 2020). Within each human TF or vTR, predictions on 53-aa tiles at 1-aa resolution were generated using PADDLE and smoothed with a 9-aa moving average. High-strength ADs were defined as the union of at least 5 consecutive 53-aa tiles with a predicted Z-score greater than 6. ADs defined in this way that overlapped by more than 26 aa were combined. Medium-strength ADs were defined similarly with a Z-score cutoff of 4 and were required to not overlap with high-strength ADs by more than 26 aa. All protein positions reported in the text are 1-indexed and inclusive; all annotations reported in supplemental tables are 0-indexed and inclusive of the start position but exclusive of the stop position. For testing in a luciferase assay, 50 predicted high-strength ADs from human TFs were chosen at random and the tile with the strongest (smoothed) predicted Z-score from each AD was used.

Yeast nuclear proteins were defined by Gene Ontology term GO:0005634 (Nucleus). Within each protein, predictions on 53-aa tiles at 1-aa resolution were generated using PADDLE and smoothed with a 9-aa moving average. All predicted Z-score local maxima higher than 3.09 ($P < 0.001$) were considered for experimental testing. The 53-aa tiles corresponding to the local maxima were included, starting from those with the largest predicted Z-score, unless they overlapped a previously-included tile by at least 15 aa. Analysis of experimentally-confirmed ADs focused on proteins that had experimental or high-throughput evidence for nuclear localization. Proteins involved in transcriptional regulation were defined by GO:0006355 (regulation of transcription, DNA-templated). To calculate enrichment of activating tiles in coactivator proteins, we generated a null model in which “activating” tiles, equal in number to actual activating tiles discovered in our screen, were randomly selected from among non-TF nuclear proteins. We then counted how many of these randomly-selected tiles were in coactivator proteins, and then repeated this sampling 10,000 times to generate a null distribution. The actual number of tiles, 42, is 2.9-fold more than the mean of the null; $P < 10^{-12}$ by one tailed Z-test. Predicted disorder for each tile was computed from D2P2 annotations as the fraction of the tile’s residues that were annotated as disordered among all 8 of the predictors listed in D2P2.

To define core ADs, predictions of sequences shorter than 53 aa were done using PADDLE-noSS by embedding the shorter region in 100 different neutral sequences composed of AGSTNQV residues and averaging the predicted values. Predictions of all tiles 5-30 aa in length within every AD were computed in this manner, and used to define core ADs for Figure 3F-G, and 7A-B. Specifically, core ADs were defined as the shortest tiles with predicted Z-score greater than 3.09 ($P < 0.001$) that do not overlap another shorter core AD. In Figure 3H, the 20-aa region within each AD with the highest predicted Z-score was included if it had Z-score greater than 3.09, and the effects of single AA mutants in these 20-aa core ADs were predicted in the same manner using PADDLE-noSS.

**Computational Peptide Docking using FlexPepDock**

To generate structural models of binding, we took 28 core ADs peptides 13 residues in length and docked them to the ABDs using Rosetta3 (Leaver-Fay et al., 2011), specifically its FlexPepDock *ab-initio* pipeline (Raveh et al., 2011). Core ADs were used as the peptide, and activator-binding domain as the receptor. Peptides were initialized in an extended conformation using Rosetta’s BuildPeptide application and manually placed near the binding pocket. For the KIX domain, NMR chemical shift data was used to aid in placement (Thakur et al., 2008). For ABD1, the AD of Gcn4 was used to aid in placement. For all models of ABD1, the first conformer from PDB structure 2LPB was used (Brzovic et al., 2011).

For each AD-ABD pair, a set of 50,000 structural models were generated and ranked by FlexPepDock’s reweighted_sc score. Binding poses were generated by clustering the top 500 ranked models using Rosetta’s cluster application with the default clustering distance threshold of 2 Å. The best-scoring model from each cluster was designated as the cluster representative, and the cluster representatives that were also among the 10 best-scoring models were used for further analysis. Additionally, larger runs with 500,000 structural models and clustering on the top 5000 models were run for 6 individual ADs for the KIX ABD. The full list of ABDs and ADs used is available in Table S5. Due to the large number of peptides that needed to be docked, the FlexPepDock *ab-initio* pipeline was modified to run in a cluster setting at a large scale and used approximately 100,000 CPU hours on the Stanford Sherlock academic cluster.

Secondary structure was calculated using DSSP (Kabsch and Sander, 1983; Touw et al., 2015) and annotations of G, H, or I were considered alpha-helical. Interaction types were computed with GetContacts (Venkatakrishnan et al., 2019) and per-AA accessible surface areas were computed with FreeSASA (Mitternacht, 2016). For each final docked structural model, buried peptide surface area was taken by subtracting the accessible surface area of the peptide in complex with the receptor from the accessible surface area of the peptide on its own. For Figure 6F, marked positions correspond to atom CG of Asp, atom CG of Leu, and the midpoint of the CG and CZ atoms in Phe (the center of the aromatic ring). Residues are shown from all the 10 best models from different clusters for all core AD structures if the marked position is within 3 Angstroms of the KIX domain or ABD1 structure.

Adapted FlexPepDock code, and wrapper code for GetContacts and FreeSASA are available at https://github.com/drorlab/med15.
